# Contribution of parasite and host genotype to immunopathology of schistosome infections

**DOI:** 10.1101/2024.01.12.574230

**Authors:** Kathrin S. Jutzeler, Winka Le Clec’h, Frédéric D. Chevalier, Timothy J.C. Anderson

## Abstract

**Background:** The role of pathogen genotype in determining disease severity and immunopathology has been studied intensively in microbial pathogens including bacteria, fungi, protozoa, and viruses, but is poorly understood in parasitic helminths. The medically important blood fluke *Schistosoma mansoni* is an excellent model system to study the impact of helminth genetic variation on immunopathology. Our laboratory has demonstrated that laboratory schistosome populations differ in sporocyst growth and cercarial production in the intermediate snail host and worm establishment and fecundity in the vertebrate host. Here, we (i) investigate the hypothesis that schistosome genotype plays a significant role in immunopathology and related parasite life history traits in the vertebrate mouse host and (ii) quantify the relative impact of parasite and host genetics on infection outcomes.

**Methods:** We infected BALB/c and C57BL/6 mice with four different laboratory schistosome populations from Africa and the Americas. We quantified disease progression in the vertebrate host by measuring body weight and complete blood count (CBC) with differential over an infection period of 12 weeks. On sacrifice, we assessed parasitological (egg and worm counts, fecundity), immunopathological (organ measurements and histopathology), and immunological (CBC with differential and cytokine profiles) characteristics to determine the impact of parasite and host genetics.

**Results:** We found significant variation between parasite populations in worm numbers, fecundity, liver and intestine egg counts, liver and spleen weight, and fibrotic area, but not in granuloma size. Variation in organ weight was explained by egg burden and by intrinsic parasite factors independent of egg burden. We found significant variation between infected mouse lines in cytokines (IFN-γ, TNF-α), eosinophil, lymphocyte, and monocyte counts.

**Conclusions:** This study showed that both parasite and host genotype impact the outcome of infection. While host genotype explains most of the variation in immunological traits, parasite genotype explains most of the variation in parasitological traits, and both host and parasite genotype impact immunopathology outcomes.

## Background

Pathogen genetic variation can play a crucial role in shaping immunopathology within the host. The recent COVID-19 pandemic serves as a pertinent example, as the emergence of new variants of SARS-CoV-2 led to changes in pathogenicity, immunity, and symptomatology in infected individuals [1–3]. Similar observations have been made in the fields of bacteriology and parasitology. *Escherichia coli* genotypes exhibit a spectrum of disease severity ranging from asymptomatic to severe, and parasite genotype explains 83% of the variation in mortality from visceral leishmaniasis [4–6]. Yet we know little about the impact of helminth genetic variation on disease severity. Helminths infect approximately 25% of the world’s human population and are ubiquitous parasites of vertebrates, invertebrates and plants. In specific cases, such as *Schistocephalus solidus* tapeworms of sticklebacks, host immune gene expression and immunological parameters vary depending on the parasite population they are infected with [7]. Moreover, genetic variation impacts infection rate, egg burden per cyst, and egg hatching rate in the sugarbeet nematode *Heterodera schachtii*, with implications for pathogenicity in plants [8]. Different *Trichinella spiralis* nematode isolates exhibit significant variation in infection clearance, larval burden, and host-inflammatory response in mammalian hosts [9,10]. These examples suggest that parasite genetic variation may have a profound impact on host responses and pathogenicity in helminth infections across diverse species.

*Schistosoma mansoni*, a blood fluke prevalent in Africa, the Caribbean, and South America, causes significant morbidity and mortality in infected people as a result of a vigorous immune response to schistosome eggs and granuloma formation around eggs that become lodged in ectopic tissues, causing inflammation, fibrosis, and portal hypertension [11]. This parasite provides a tractable model organism for investigating the influence of helminth genetic variation on immunopathology, because it can be maintained in the laboratory using rodents as definitive hosts and aquatic snails as intermediate hosts. Schistosomes vary in multiple heritable traits including drug resistance and snail host specificity [12–15]. We have previously demonstrated dramatic differences in cercarial shedding numbers and mortality of intermediate snail hosts infected with two different parasite populations from Brazil, SmBRE or SmLE [16]. Low or high shedding populations derived from crosses between these two parasite populations also varied in life history traits in the rodent host. Low shedding parasites showed lower fecundity, reduced hepatosplenomegaly and hepatic fibrosis in mice than high shedding parasites [17].

The striking differences between these two populations led us to hypothesize that parasite genotype may influence other life history traits and immunopathology in the vertebrate mouse host. Several studies support this notion: Anderson and Cheever [18] first documented significant differences in worm and egg burden when mice were infected with *S. mansoni* field isolates obtained from different geographical regions. Subsequent studies utilizing laboratory-maintained schistosome populations have further examined various parameters including egg dimensions and granuloma burden/area in the liver and intestine (Table 1) [19–21].

**Table 1.**
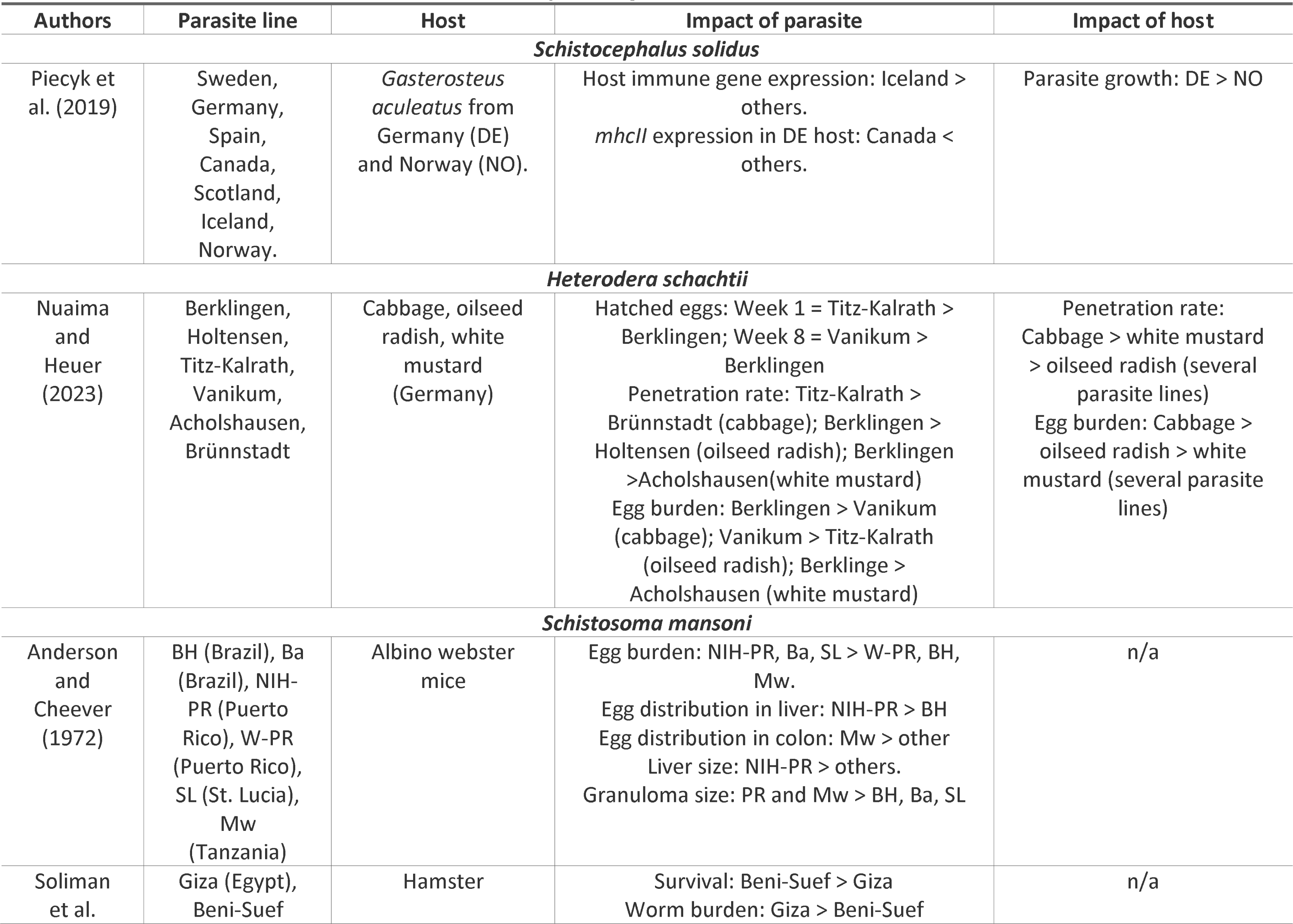
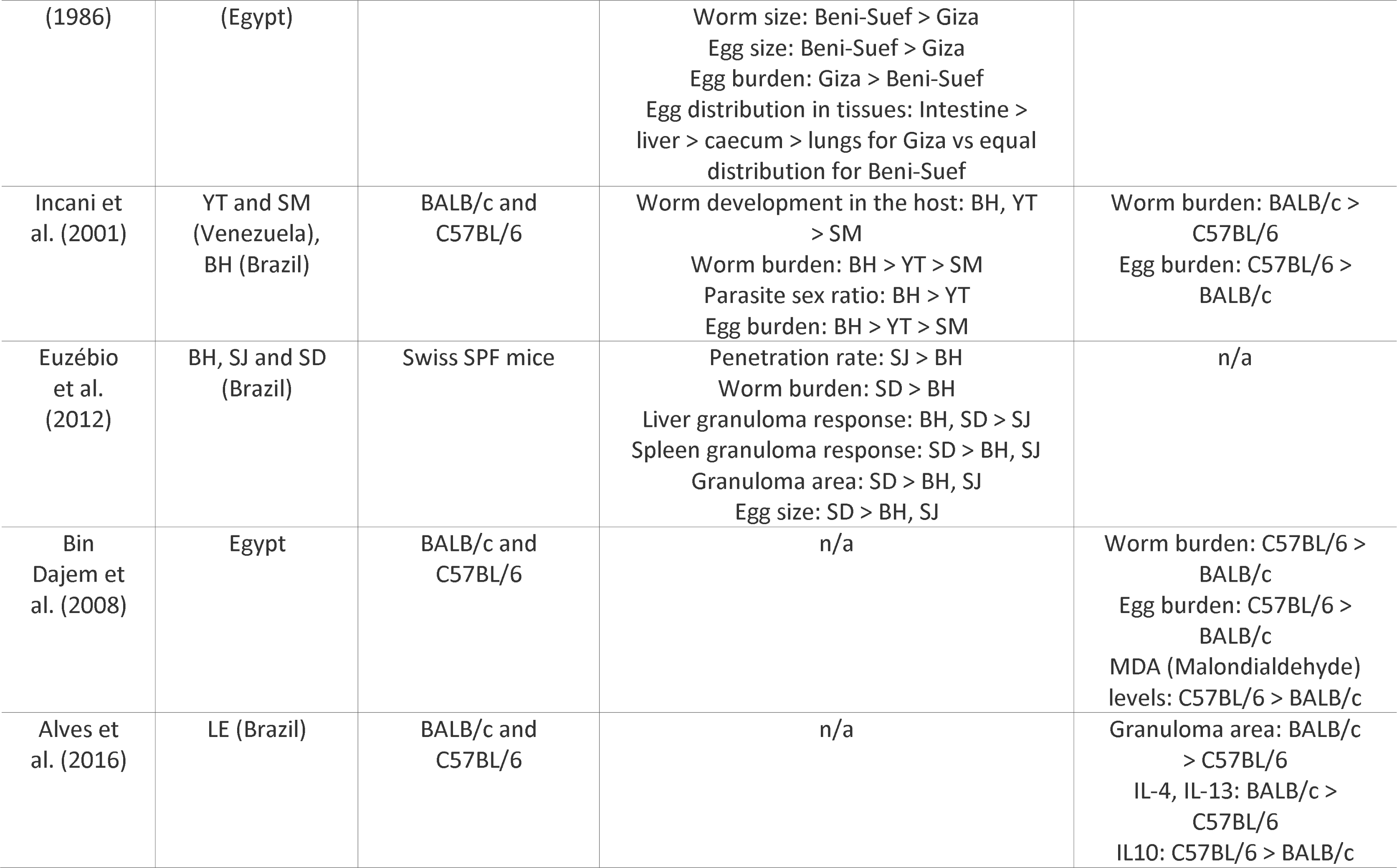
Previously Investigated Parasite and Host Traits.

We maintain four distinct *S. mansoni* populations in our laboratory (SmEG from Egypt, SmLE and SmBRE from Brazil, and SmOR [oxamniquine resistant], a cross between SmLE and SmHR from Puerto Rico) which we used in this study. Our central aim was to investigate the impact of parasite genetics on immunopathology while accounting for host genetic differences. To do this, we infected mice with schistosome larvae from SmBRE, SmLE, SmEG, or SmOR parasite populations. Over the course of the infection, we monitored body weight and conducted complete blood counts. On sacrifice, we measured organ weights, assessed hepatic fibrosis and granuloma size, and analyzed cytokine levels to evaluate the disease progression and immunopathological changes induced by each parasite population. Host genetic background may also influence parasite-induced immunopathology. We therefore conducted these experiments using both BALB/c and C57BL/6 inbred lines. These mouse strains are commonly used in laboratory studies and differ in their innate and adaptive immune responses. BALB/c mice mount a dominant Th2 immune response to infections, while C57BL/6 mice display a Th1 bias [22,23]. Naïve BALB/c CD4+ T cells produce more IL-4 than CD4+ T cells from C57BL/6 mice, which induces the proliferation of CD4+ Th2 cells, and lower levels of the pro-inflammatory cytokine IL-12 [24,25].

Our results revealed that parasite genetics explained most variation in parasite traits, while immunological traits were most strongly influenced by the host. However, both parasite and host genetics affected immunopathology traits. These results provide compelling evidence for the role of parasite genetics in modulating *S. mansoni*-induced immunopathology in mouse hosts.

## Methods

### Ethics statement

This study was performed in accordance with the Guide for the Care and Use of Laboratory Animals of the National Institutes of Health. The protocol was approved by the Institutional Animal Care and Use Committee of Texas Biomedical Research Institute (permit number: 1420-MU).

### Overview of study design

Our study design is summarized in Figure 1, and the methodology for each stage is explained below.

**Figure 1:**
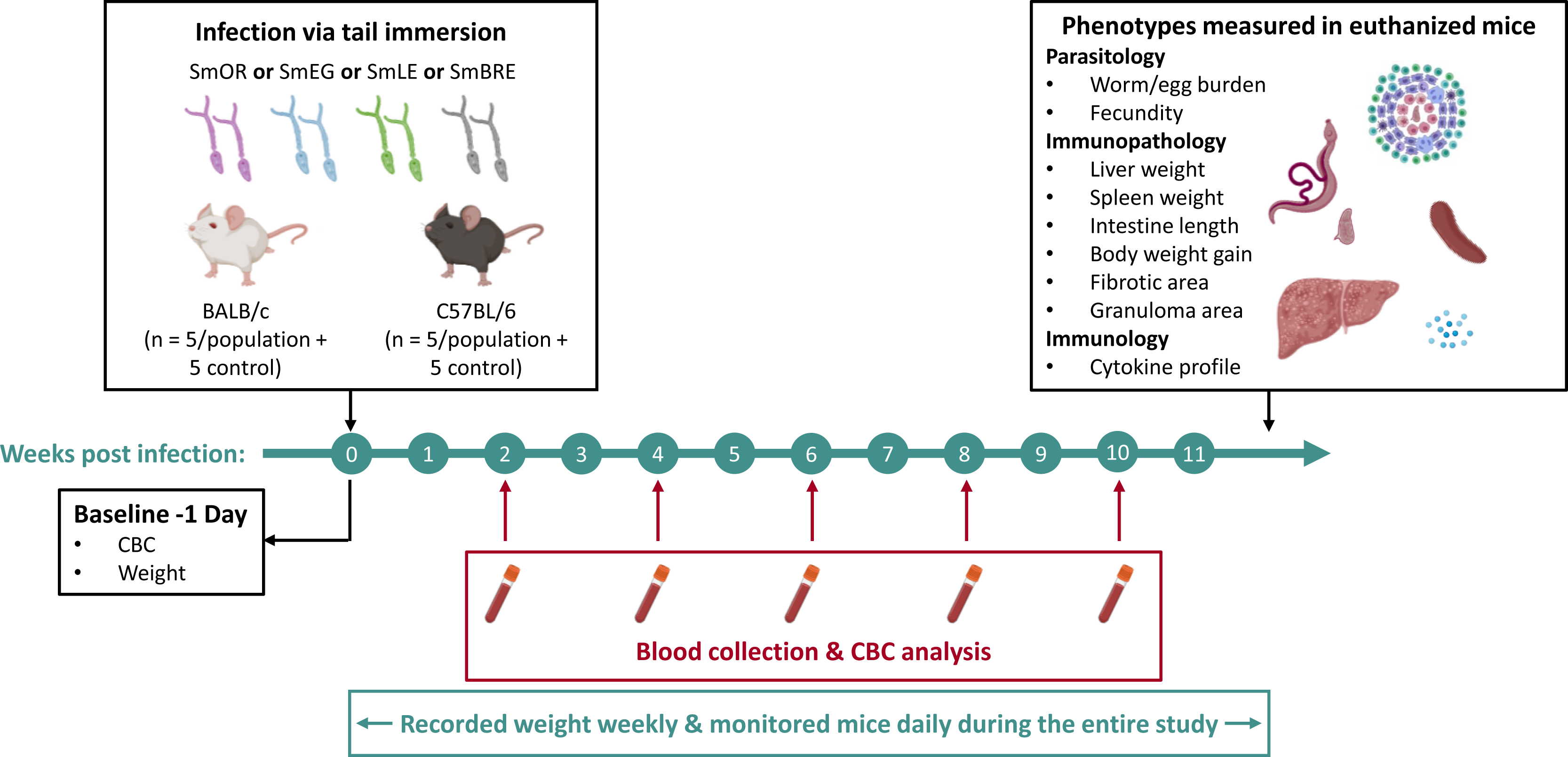
Experimental timeline. We investigated the influence of parasite and host genotype on disease progression during schistosome infection in BALB/c and C57BL/6 mice. Four laboratory schistosome populations from Africa (SmEG) and the Americas (SmOR, SmLE, and SmBRE) were used to infect the mice. Over a 12-week infection period, we quantified disease progression in the vertebrate host by monitoring body weight and complete blood count (CBC). Upon sacrifice, we measured multiple parasitological, immunopathological, and immunological traits (see box, top right and main text).

#### *Schistosoma mansoni* parasites and mouse infection

We used SmLE (from Belo Horizonte, Brazil), SmBRE (from Recife, Brazil), SmEG (from the Theodor Bilharz Research Institute, Cairo, Egypt) and SmOR (oxamniquine resistant population homozygous for the deletion in amino acid 142 in *SmSULT-OR*) parasite lines in this study. We placed 10-20 *Biomphalaria glabrata* (Bg36 for SmLE and SmOR, BgBRE for SmBRE) and *B. alexandrina* (for SmEG) snails in beakers and shed them in artificial pond water for 2 hours under light. We infected 5 female BALB/c and 5 female C57BL/6 mice (Envigo, 7-9 weeks old) per parasite population with 50 cercariae via tail immersion [26]. Control cohorts (N = 5 per mouse line) were mock infected by placing tails in glass vials containing water but no cercariae. We conducted these infections over a period of 12 days to accommodate all 50 mice, but randomized cage allocation to avoid batch effects (Fig. 1).

### Measuring parasitological, immunopathological, and immunological traits during infection and upon sacrifice

We collected phenotypic data from each mouse during the infection period and upon euthanasia 12 weeks after the infection. We categorized each trait as parasitological, immunopathological, or immunological:

#### Parasitological traits

##### Penetration rate

We counted whole cercariae and heads that remained in the glass vial after mouse infection to calculate the penetration rate:

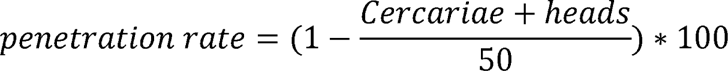

##### Worm and egg counts in the liver and the intestine

We perfused each mouse to collect and count worms by sex [27]. To count liver eggs, the median and caudate lobes were weighed and stored in a 50ml conical tube. The intestine was detached from the mesentery and the anus and flushed with a 1x PBS solution. We withdrew as much liquid as possible before weighing the organ and placing it in a 15 ml conical tube. Both tubes were filled with 4% KOH and stored overnight at 37°C (without agitation). The next day, we centrifuged the tubes (1500 rpm for 5 minutes) and removed half of the supernatant. We counted eggs in both organs in triplicate in 50-100 µl of tissue suspension and calculated the number of eggs per g of tissue as follows:

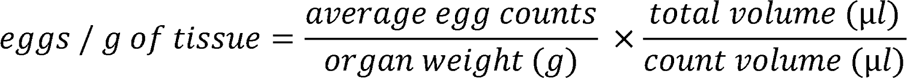

##### Fecundity

We calculated fecundity as the total number of eggs divided by the number of female worms recovered after perfusion.

#### Immunopathological traits

##### Body weight

We weighed each mouse on the day of infection and weekly thereafter on a Mettler Toledo MS12002TS scale. We calculated the percent weight change as follows:

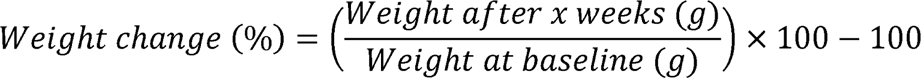

##### Liver spleen and spleen weight

We weighed each liver and spleen on a Mettler Toledo PB3002-S scale. We normalized the organ weight to account for body weight as follows:

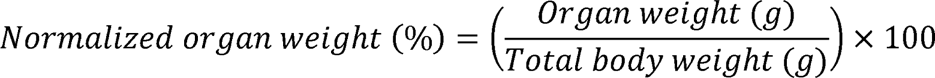

##### Intestine length

Each intestine was removed and photographed. We used the polygon tool from ImageJ 1.53k [28] to trace the intestine and return the intestine length with the built-in *Measure* command.

##### Liver histology to assess granuloma and fibrotic area

We stored the left frontal lobe of each liver in 10% neutral buffered formalin (N=47 liver samples) for further processing and embedding into paraffin. Each tissue block was cut into 4 µm sections using a Microm HM325 rotary microtome. We cut sections at 60 µm and mounted 12 sections per liver onto four slides. The slides were alternately stained with hematoxylin-eosin for quantitative analysis of granulomas or Masson’s trichrome to detect fibrosis due to collagen deposition [29]. All slides were scanned with the Zeiss Axio Scan.Z1 whole slide scanner at a resolution of 0.22 µm/pixel. We analyzed sections that were at least 120 µm apart using HALO® software (v3.4, Indica labs). We annotated and quantified the area of individual granulomas (16-44/liver) surrounding single schistosome eggs, recording 118-152 granulomas for each experimental group. To assess fibrotic area, we trained HALO’s Area Quantification tool (v2.3.1) on the different trichrome dyes (blue, red, brown). This tool automatically detects pixels of an assigned color and calculates the total area stained with each dye.

#### Immunological traits

##### Complete Blood Count (CBC) with differential

We obtained CBC profiles with differential to measure the number of each white blood cell type from individual mice before infection and then bi-weekly until week 10. Briefly, we used a 4 or 5-mm lancet (Goldenrod Animal Lancet, Medipoint, Mineola, NY) to puncture the submandibular vein and collected the blood in an EDTA tube (Microtainer, Becton Dickenson). We injected mice with 50-100 µl 5% saline solution to replace fluids and promote recovery. Blood was analyzed using a ProCyte Dx Hematology Analyzer (IDEXX Laboratories, Inc., Westbrook, ME).

##### Cytokine assay

We collected 100-200 µl of blood by cardiac puncture after euthanasia but before perfusion. We stored the blood in a microcentrifuge tube for 20-23 minutes at room temperature before spinning the tube at 2,000 × g for 10 minutes. We collected the serum fraction in a new tube and placed it on dry ice until permanent storage at –80°C. We used the LEGENDplex™ Mouse Th1/Th2 Cytokine Panel V03 (BioLegend®, San Diego, CA) to simultaneously quantify eight cytokines (IFN-γ, IL-5, TNF-α, IL-2, IL-6, IL-4, IL-10, and IL-13) secreted by Th1 and/or Th2 cells. Following the manufacturer instructions, we diluted 12.5 μl of serum with 12.5 μl of assay buffer and prepared reactions in 96-well V-bottom plates. We transferred the reactions to sterile snap cap tubes (VWR) and read the samples on a BD Symphony Analyzer. The analysis was conducted using LEGENDplex™ Cloud-based Data Analysis Software (https://legendplex.qognit.com/).

### Statistical analysis

#### Trait-by-trait analyses

We performed all statistical analyses and plotted graphs using R software (v4.2.0) and package *rstatix v0.7.2* [30,31]. We excluded the control group from statistical analyses examining differences between parasite populations in immunopathology. For normally distributed data (Shapiro test, *p* > 0.05) with homogenous variance (Bartlett test, *p* > 0.05), we used one-way ANOVA followed by the Tukey HSD post-hoc test. For non-normally distributed data, we applied the Kruskal-Wallis test followed by Dunn’s post-hoc test or the Friedman test followed by Conover’s post-hoc test for longitudinal analysis. We adjusted *p*-values for multiple comparisons using the Benjamini–Hochberg method and considered these significant when *p* < 0.05 [32]. To make comparisons between hosts, we performed Wilcoxon rank-sum tests (non-parametric) or Student’s t-tests (parametric).

#### General linear modeling

We suspected that some differences in immunopathology may be explained by differences in egg counts among parasite populations. We conducted GLM of all phenotypes using the function glm() from the *stats* R package. We assessed the fit of linear models by examining the distribution of residuals, quantile-quantile, scale-location plots, and Bayesian information criterion (BIC) and normalized the dependent variable with log (liver weight, fibrosis, intestine length, granuloma area, all cytokine data, lymphocytes, reticulocytes) or square root (spleen weight, eosinophils, neutrophils, monocytes) transformation to improve the fit of the model when applicable [33]. The following model showed the best fit for all parasitological, immunopathological, and cytokine measures except liver weight:

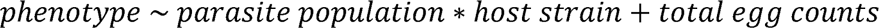

We used an interaction model exclusively for liver weight:

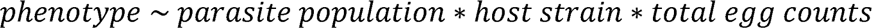

To account for baseline values, we used the following model for CBC data:

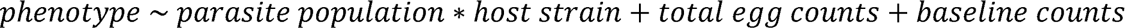

#### Contribution of parasite and host to immunopathology

To determine effect size of parasite population and host genotype, we first log-transformed non-normally distributed data and then used the R function aov() from the *stats* package to perform an ANOVA test for each parameter before calculating effect sizes with the eta_squared() function from the *effectsize v0.8.6* package for R [34,35].

## Results

### Parasitological phenotypes

#### Penetration rates

We measured parasite penetration rates by counting whole cercariae and heads that remained in the glass vial after mouse infection [36]. We found no significant differences between experimental groups (Fig. 2A; Kruskal-Wallis; BALB/c: H = 4.15, df = 3, *P* = 0.246; C57BL/6: H 2.5, df = 3, *P* = 0.475; Wilcoxon test to compare hosts; BRE: W = 13, *P* = 1, EG: W = 11.5, *P* = 1, LE: W = 6.5, *P* = 0.984, OR: W = 9, *P* = 1).

**Figure 2:**
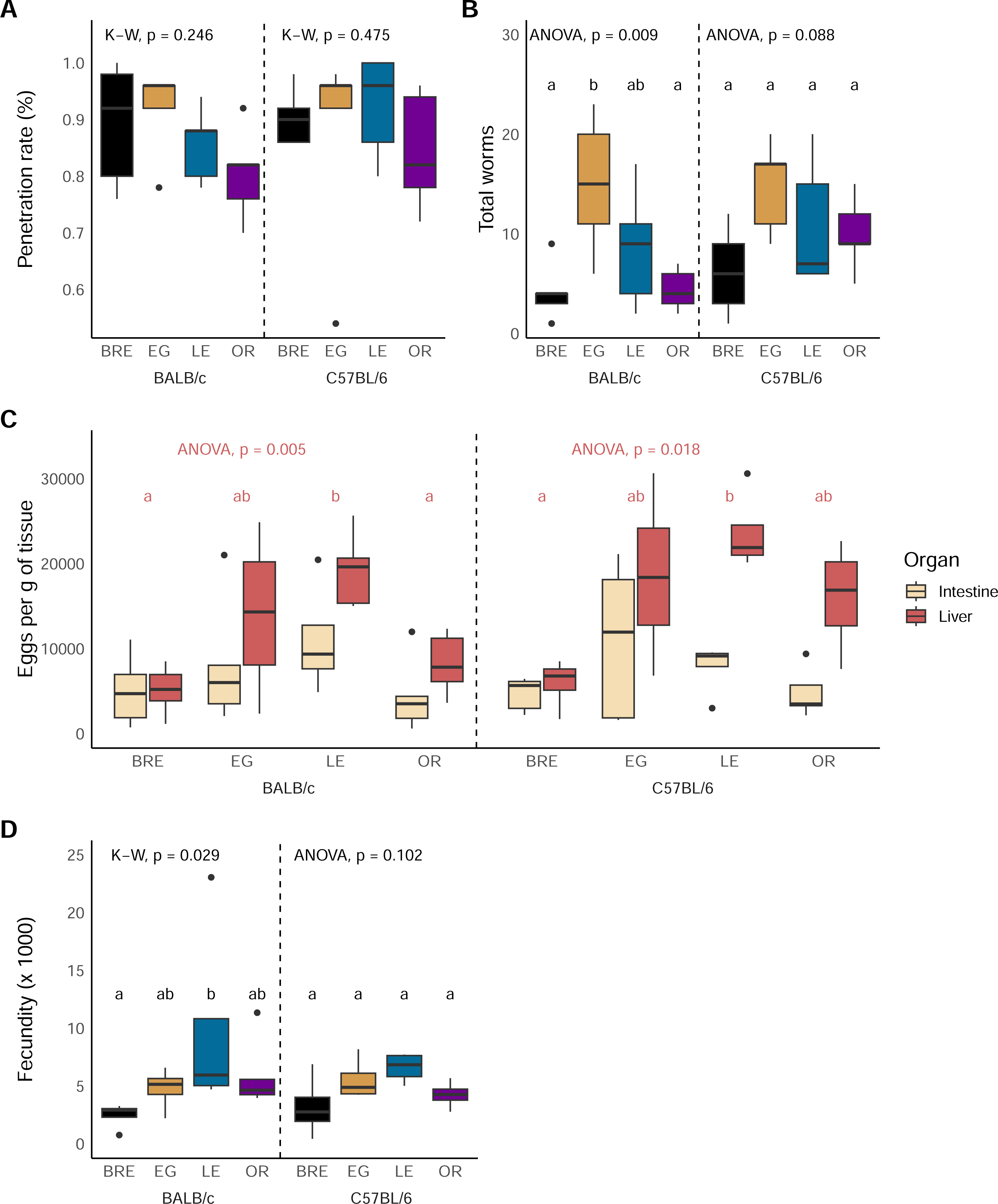
Parasitological outcome is driven by parasite rather than host genotype. Box and whisker plots showing cercarial penetration rate **(A)**, worm burden **(B)**, liver and intestine egg counts **(C)** and fecundity **(D)** for infections with SmBRE, SmEG, SmLE and SmOR in BALB/c (left) and C57BL/6 mice (right). Parasite populations marked with different letters are significantly different in comparisons using Kruskal-Wallis (K-W) followed by Dunn’s or ANOVA followed by Tukey HSD post hoc test. Use of ANOVA or Kruskal-Wallis is shown at the top of each graph. For (C), stats are shown for liver eggs only.

#### Host survival

All C57BL/6 mice survived throughout the study period. However, we sacrificed three BALB/c mice due to severe disease progression during the study (SmLE: days 54 and 74, SmEG: day 83), and two BALB/c mice died from the infection without prior symptoms (SmEG: day 63, SmOR: day 75).

#### Worm and egg burden

On perfusion, we observed significant variation in worm burden among BALB/c mice and almost significant variation for C57BL/6 mice infected with the four parasite populations (Fig. 2B; ANOVA; BALB/c: F_(3, 16)_ = 5.4, *P* = 0.009; C57BL/6: F_(3, 16)_ = 2.6, *P* = 0.088). SmEG caused significantly higher worm burden than SmBRE and SmOR in BALB/c mice. Liver egg burden varied significantly in both BALB/c and C57BL/6 mice infected with the four parasite populations (Fig. 2C; ANOVA; BALB/c: F_(3, 15)_ = 6.44, *P* = 0.005; C57BL/6: F_(3, 12)_ = 4.99, *P* = 0.018). SmLE infected mice had the highest liver egg burden, while SmBRE caused the lowest burden in both mouse strains. Comparisons between host strains revealed no significant differences in terms of parasite and egg burden.

#### Fecundity

Fecundity was significantly higher in BALB/c mice that were infected with SmLE compared to SmBRE (Fig. 2D; Kruskal-Wallis, H = 9.02, df = 3, *P* = 0.0291). We did not detect an impact of host genotype on worm fecundity.

#### Tissue distribution of eggs

Visual inspection of Fig. 2C suggests differences in tissue distribution of eggs for the four parasite populations. We therefore examined tissue tropism by comparing egg burden in liver and gut. However, the distribution of eggs in SmEG, SmLE, SmOR and SmBRE-infected mice was similar in both tissues, and tissue distribution was not influenced by host genotype (Fig. 2C). We conducted a separate analysis comparing the ratio between liver and intestine eggs in individual mice, but this similarly revealed no significant differences (Additional file 1: Fig. S1).

### Parasite genotype and parasite burden impact tissue damage

Next, we analyzed immunopathological phenotypes in the infected mice. To focus on differences between parasite populations rather than host infection status, we excluded the control group from the following statistical analyses.

#### Mouse growth

Schistosome-infected mice typically experience weight gain during the early stages of the infection [37–39]. Schistosome eggs are usually excreted through the intestine, but can migrate to other tissues and induce hepatosplenomegaly in both humans and mice [40], which may contribute to the observed increase in body weight. All mice showed continuous weight gain over the course of the study. However, SmLE infected BALB/c mice exhibited a significantly reduced weight gain compared to BALB/c mice infected with other parasite populations (Fig. 3A; Friedman test; BALB/c: χ2 = 18.7, df = 3, *P* < 0.001). We did not detect any significant differences between the infected C57BL/6 mice (Fig. 3A; Friedman test; C57BL/6: χ2 = 1.7, df = 3, *P* = 0.637).

**Figure 3:**
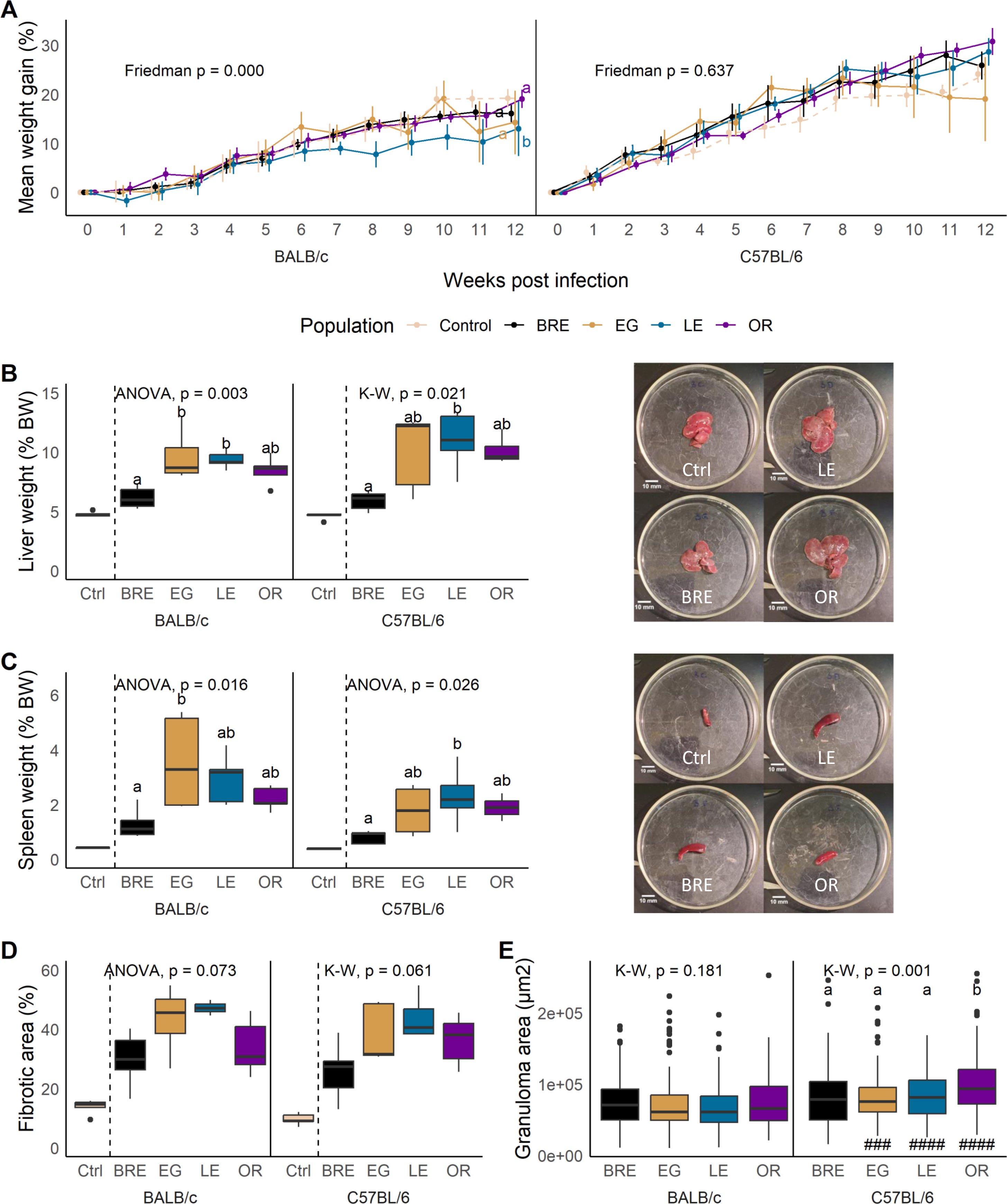
Impact of parasite and host on mouse immunopathology. **(A)** Longitudinal plots of weight gain in mice infected with the 4 parasite populations (solid lines) and uninfected controls (dotted lines). Means shown with standard error. **(B)** Liver weight or **(C)** spleen weight from euthanized BALB/c (left) and C57BL/6 mice (right) infected with the 4 parasite populations or uninfected controls. Photos (right) show representative organ samples from mice infected with 3 of the 4 parasite populations and the control group for reference. **(D)** Fibrotic area and **(E)** granuloma area measured from liver sections. For all box plots, statistical comparisons are between infected groups only. Parasite populations marked with different letters are significantly different in comparisons using Kruskal-Wallis (K-W) followed by Dunn’s or ANOVA followed by Tukey HSD post hoc test. Differences between BALB/c (left) and C57BL/6 mice (right) in the pathology traits were calculated with Wilcoxon rank-sum or Student’s t-test and are shown by: # *P* < 0.05, ## *P* < 0.01, ### *P* < 0.001.

#### Liver and spleen weight and intestine length

All infected mice exhibited hepatosplenomegaly compared to uninfected control groups. Livers of SmBRE infected mice were significantly smaller than livers of SmLE infected mice (regardless of the host strain) and SmEG infected BALB/c mice (Fig. 3B; BALB/c: ANOVA, F_(3, 16)_ = 7.27, *P =* 0.003; C57BL/6: Kruskal-Wallis, H = 9.79, df = 3, *P* = 0.0205). Similarly, mice infected with SmBRE had reduced spleen mass compared to mice infected with other parasite populations. Notably, SmEG-infected BALB/c and SmLE-infected C57BL/6 mice showed a significant increase in spleen weight (Fig. 3C; ANOVA; BALB/c: F_(3, 16)_ = 4.68, *P =* 0.016; C57BL/6: F_(3, 16)_ = 4.01, *P =* 0.026). Intestine length was significantly shorter in C57BL/6 compared to BALB/c mice infected with SmLE (Additional file 2: Fig. S2).

#### Fibrotic area and granuloma size

Fibrotic area showed the same trend as liver and spleen weight, though the differences were not statistically significant. All infected mice had an increase of fibrotic liver tissue, with SmBRE causing less fibrosis than other parasite populations (Fig. 3D; BALB/c: ANOVA, F_(3, 13)_ = 2.94, *P* = 0.073; C57BL/6: Kruskal-Wallis, H = 7.41, df = 3, *P* = 0.0599). Contrary to other reports that noted increased granuloma area in BALB/c mice [41,42], we measured larger granulomas in infected C57BL/6 mice (Fig. 3E; Wilcoxon test; EG: W = 5383, *P* < 0.001; LE: W = 5212, *P* < 0.001, OR: W = 5391, *P* < 0.001). In addition, infection of C57BL/6 mice with SmOR resulted in the formation of larger granulomas compared to other parasite populations (Fig. 3E; Kruskal-Wallis; BALB/c: H = 4.88, df = 3, *P* = 0.181; C57BL/6: H = 16.2, df = 3, *P* = 0.00105).

#### Does parasite genotype and/or egg burden impact organ pathology?

We used generalized linear modeling (GLM) to examine the effects of parasite population, host strain, total egg burden, and host-parasite interactions (Table 2). Liver weight, spleen weight, and intestine length were all heavily influenced by the total egg burden. However, we also identified a significant effect of parasite population on both liver and spleen weight. In turn, the host strain emerged as a significant factor affecting intestine length. Intriguingly, the GLM analysis unveiled a contribution approaching significance of the parasite population to fibrosis, confirming the results from the trait-by-trait analysis (Fig. 3D). We also found that host-parasite interactions significantly influenced liver and spleen weight, but not other traits examined. However, this was significant for the interaction model (liver weight) and square root transformed data (spleen weight) only.

**Table 2.**
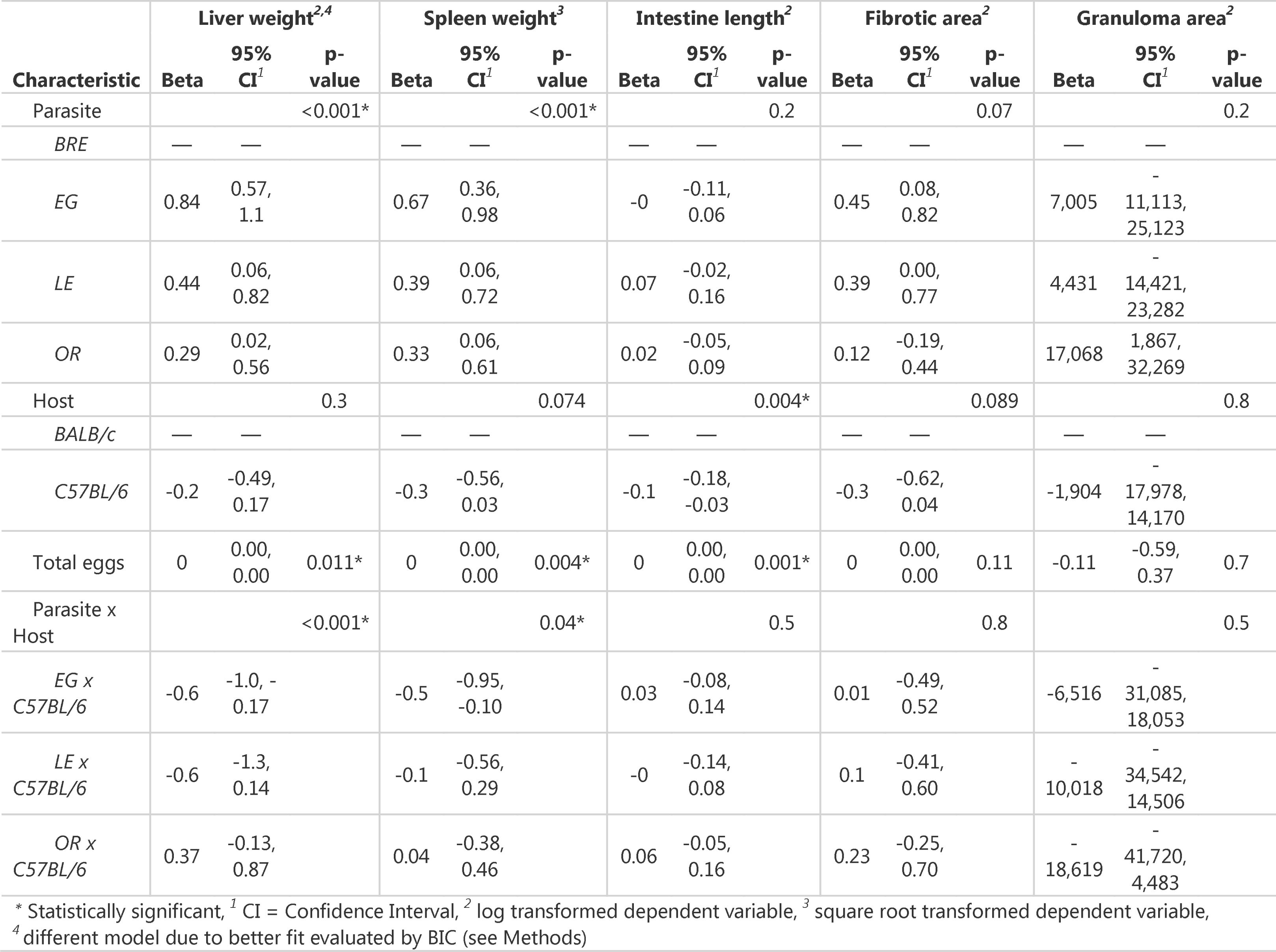
Generalized linear model output for all traits.

| Characteristic | Liver weight<sup>2,4</sup> |  |  | Spleen weight<sup>3</sup> |  |  | Intestine length<sup>2</sup> |  |  | Fibrotic area<sup>2</sup> |  |  | Granuloma area<sup>2</sup> |  |  |
| --- | --- | --- | --- | --- | --- | --- | --- | --- | --- | --- | --- | --- | --- | --- | --- |
|  | Beta | 95% CI<sup>1</sup> | p-value | Beta | 95% CI<sup>1</sup> | p-value | Beta | 95% CI<sup>1</sup> | p-value | Beta | 95% CI<sup>1</sup> | p-value | Beta | 95% CI<sup>1</sup> | p-value |
| Parasite |  |  | <0.001* |  |  | <0.001* |  |  | 0.2 |  |  | 0.07 |  |  | 0.2 |
| *BRE* | — | — |  | — | — |  | — | — |  | — | — |  | — | — |  |
| *EG* | 0.84 | 0.57, 1.1 |  | 0.67 | 0.36, 0.98 |  | -0 | -0.11, 0.06 |  | 0.45 | 0.08, 0.82 |  | 7,005 | -<br>11,113, 25,123 |  |
| *LE* | 0.44 | 0.06, 0.82 |  | 0.39 | 0.06, 0.72 |  | 0.07 | -0.02, 0.16 |  | 0.39 | 0.00, 0.77 |  | 4,431 | -<br>14,421, 23,282 |  |
| *OR* | 0.29 | 0.02, 0.56 |  | 0.33 | 0.06, 0.61 |  | 0.02 | -0.05, 0.09 |  | 0.12 | -0.19, 0.44 |  | 17,068 | 1,867, 32,269 |  |
| Host |  |  | 0.3 |  |  | 0.074 |  |  | 0.004* |  |  | 0.089 |  |  | 0.8 |
| *BALB/c* | — | — |  | — | — |  | — | — |  | — | — |  | — | — |  |
| *C57BL/6* | -0.2 | -0.49, 0.17 |  | -0.3 | -0.56, 0.03 |  | -0.1 | -0.18, -0.03 |  | -0.3 | -0.62, 0.04 |  | -1,904 | -<br>17,978, 14,170 |  |
| Total eggs | 0 | 0.00, 0.00 | 0.011* | 0 | 0.00, 0.00 | 0.004* | 0 | 0.00, 0.00 | 0.001* | 0 | 0.00, 0.00 | 0.11 | -0.11 | -0.59, 0.37 | 0.7 |
| Parasite x Host |  |  | <0.001* |  |  | 0.04* |  |  | 0.5 |  |  | 0.8 |  |  | 0.5 |
| *EG x C57BL/6* | -0.6 | -1.0, -0.17 |  | -0.5 | -0.95, -0.10 |  | 0.03 | -0.08, 0.14 |  | 0.01 | -0.49, 0.52 |  | -6,516 | -<br>31,085, 18,053 |  |
| *LE x C57BL/6* | -0.6 | -1.3, 0.14 |  | -0.1 | -0.56, 0.29 |  | -0 | -0.14, 0.08 |  | 0.1 | -0.41, 0.60 |  | -<br>10,018 | -<br>34,542, 14,506 |  |
| *OR x C57BL/6* | 0.37 | -0.13, 0.87 |  | 0.04 | -0.38, 0.46 |  | 0.06 | -0.05, 0.16 |  | 0.23 | -0.25, 0.70 |  | -<br>18,619 | -<br>41,720, 4,483 |  |

\* Statistically significant, <sup>1</sup> CI = Confidence Interval, <sup>2</sup> log transformed dependent variable, <sup>3</sup> square root transformed dependent variable,
<sup>4</sup> different model due to better fit evaluated by BIC (see Methods)

### Host genotype is associated with the immunological response to infection

#### Complete blood counts

To monitor the immune response during infection, we conducted a complete blood count (CBC) with differential every two weeks. As anticipated with helminths, eosinophilia was observed in all infected mice [43], though we did not detect any significant differences between parasite populations in either host strain (Fig. 4A, Friedman; BALB/c χ2 = 2.6, df = 3, *P =* 0.457; C57BL/6: χ2 = 5.24, df = 3, *P =* 0.155). We observed a substantial increase in basophils in SmEG infected C57BL/6 mice past the 6-week mark, yielding a significant Friedman test (Fig. 4B; Friedman; BALB/c: χ2 = 1.75, df = 3, *P =* 0.625; C57BL/6: χ2 = 8.2, df = 3, *P =* 0.042). As a consequence of *p*-value adjustment for multiple comparisons, however, the post-hoc analysis did not reveal any significant differences between parasite populations. We observed the same outcome with lymphocytes (Fig. 4C; Friedman; BALB/c: χ2 = 2, df = 3, *P =* 0.572; C57BL/6: χ2 = 10.6, df = 3, *P =* 0.014). Both monocyte (Fig. 4D) and neutrophil levels (Additional File 3: Fig. S3A) exhibited an upward trend in infected mice.

**Figure 4:**
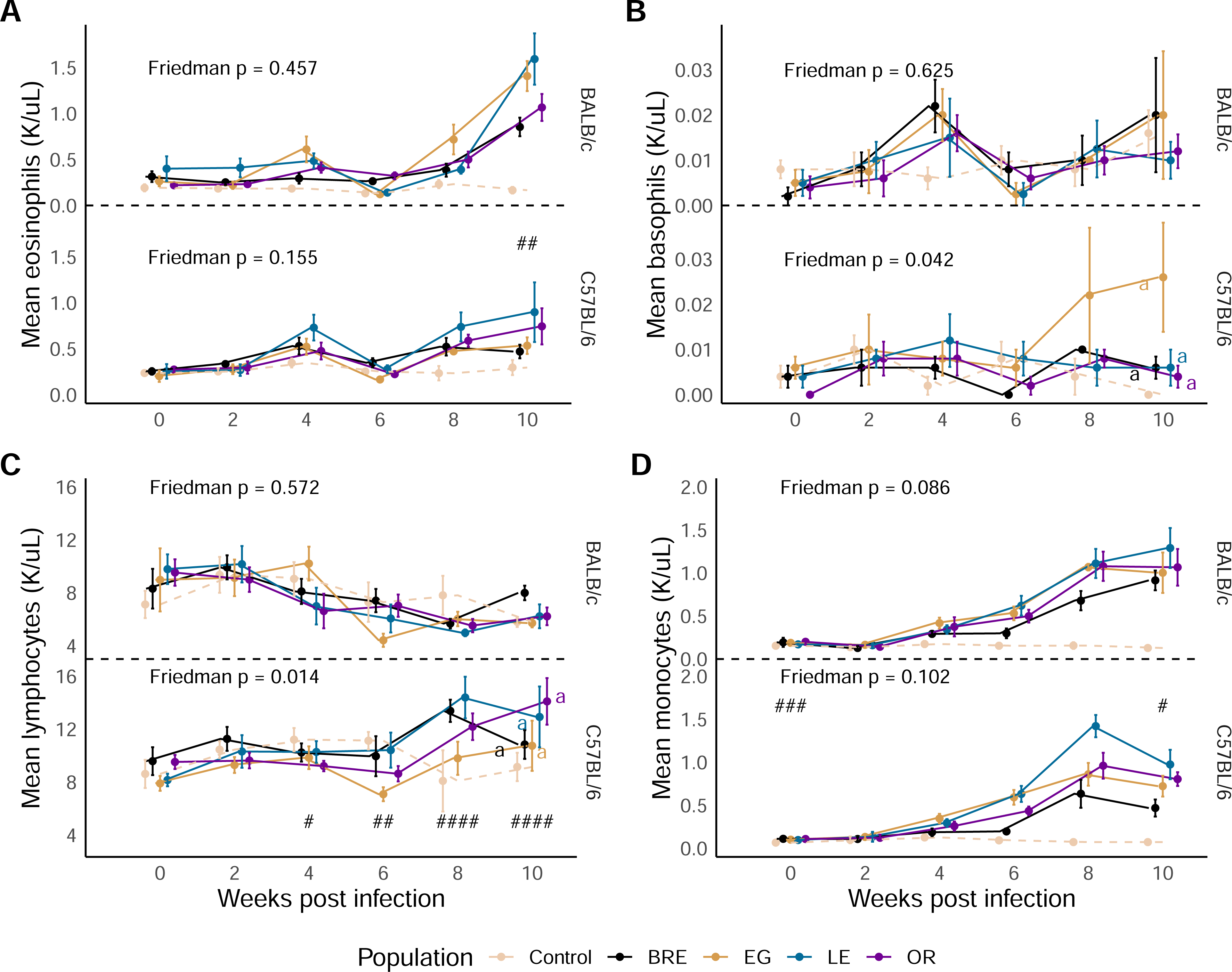
White blood cells and reticulocytes are influenced by host background. Longitudinal data of **(A)** eosinophil, **(B)** basophil, **(C)** lymphocyte and **(D)** monocyte levels in BALB/c (top) and C57BL/6 mice (bottom) infected with the 4 infected parasite populations (solid lines) or uninfected controls (dotted lines). For all plots, statistical comparisons are between infected groups only. Mice with missing data points were excluded from the analysis. Groups not connected by the same letter and separated by host strain are significantly different (Friedman’s test followed by Conover’s post hoc test). # *P* < 0.05, ## *P* < 0.01, ### *P* < 0.001: values are significantly different between infected (parasite lines combined) host strains using Wilcoxon rank-sum test.

As we did not observe significant differences between parasite populations in the CBC analysis, we grouped infected mice together to compare host strains. BALB/c mice had more circulating eosinophils in the blood after 10 weeks of infection than their C57BL/6 counterparts (Fig. 4A; Wilcoxon; Week 10: W = 305, *P* = 0.00163). In addition, lymphocyte production varied between the host strains, increasing in C57BL/6 mice after 6 weeks (Fig. 4C; Wilcoxon; Week 4: W = 91, *P* < 0.0128; Week 6: W = 71, *P* < 0.00208; Week 8: W = 1, *P* < 0.001; Week 10: W = 25, *P* < 0.001). Monocytes differed between the infected hosts at the beginning and at the end of the infection (Fig. 4C; Wilcoxon; Baseline: W = 318, *P* < 0.001; Week 10: W = 266, *P =* 0.0387), whereas neutrophil levels varied significantly between the hosts at baseline and week 4 (Additional file3: Fig. S3A).

We anticipated a decrease in hematocrit in infected animals, because schistosome parasites feed on blood [36]. Surprisingly, none of the mice exhibited signs of anemia throughout the infection period (Additional file 3: Fig. S3B), though we observed an increase in reticulocytes in the later stage of infection (Additional file 3: Fig. S3C).

We used GLM to assess contributions of the parasite, host, and egg burden, incorporating baseline values into the model for week 10 CBC measurements (Table 3). Mouse line was the dominant factor for eosinophils and monocytes, but not for lymphocyte production. In addition, total egg burden significantly impacted monocyte, basophil and reticulocyte levels. We did not see any impact of parasite genotype or host-parasite interactions.

**Table 3.**
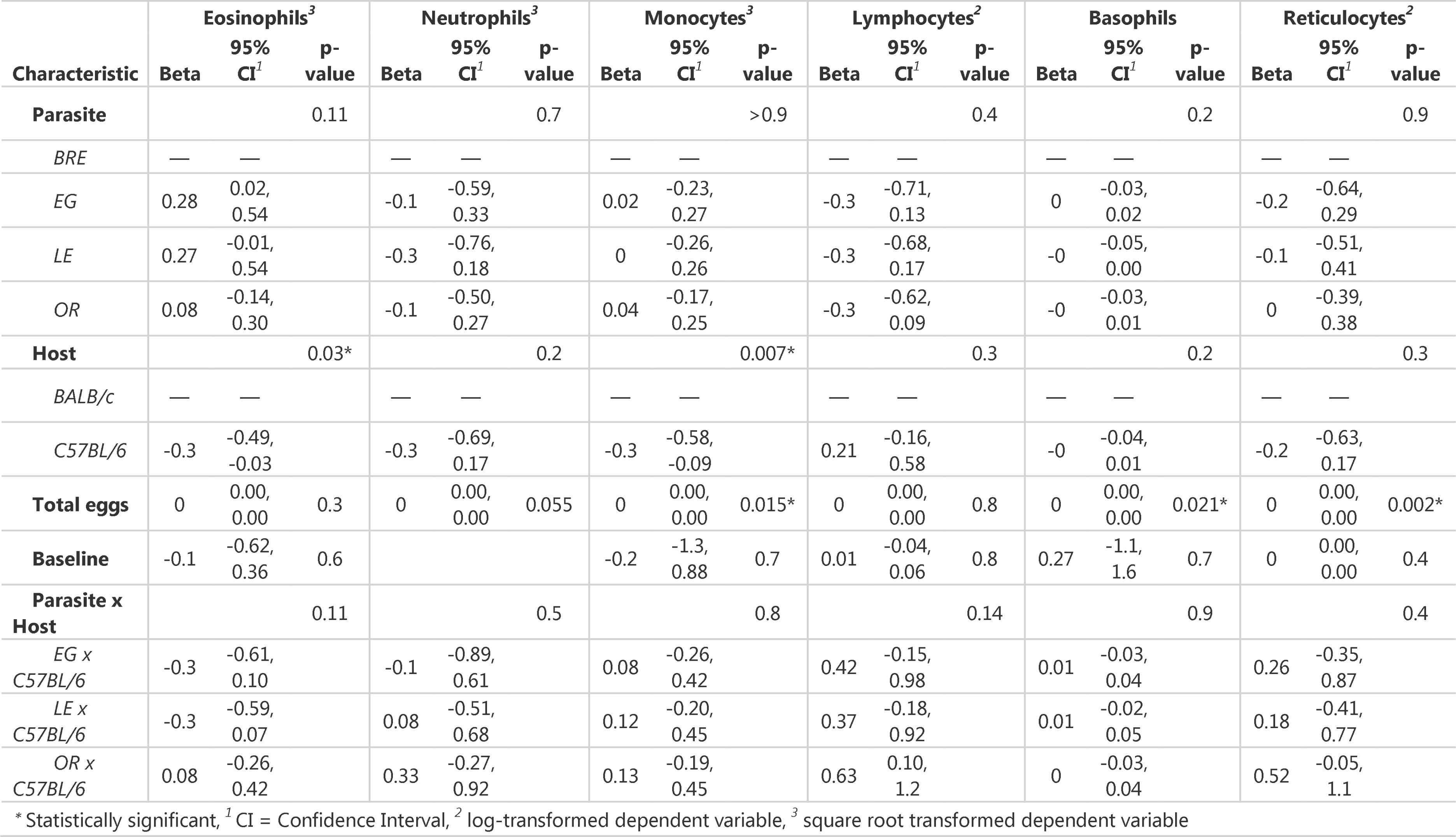
Generalized linear model output for CBC.

#### Cytokines

We quantified eight Th1/Th2 associated cytokines in serum collected at week 12. Our analysis showed no significant differences between parasite populations for any of the measured cytokines. However, we observed significant variations between the different host strains. First, IFN-γ levels were significantly elevated in BALB/c mice compared to C57BL/6 (Fig. 5A; Wilcoxon; BR, EG, LE: W = 25, *P* = 0.0106; OR: W = 20, *P* = 0.016). In contrast, C57BL/6 mice produced more TNF-α in response to schistosome infection than BALB/c mice (Fig. 5B; Wilcoxon; BRE: W = 0, *P* = 0.0318; EG: W = 6, *P* = 0.222; LE: W = 2, *P* = 0.0423, OR: W = 1, *P* = 0.0423).

**Figure 5:**
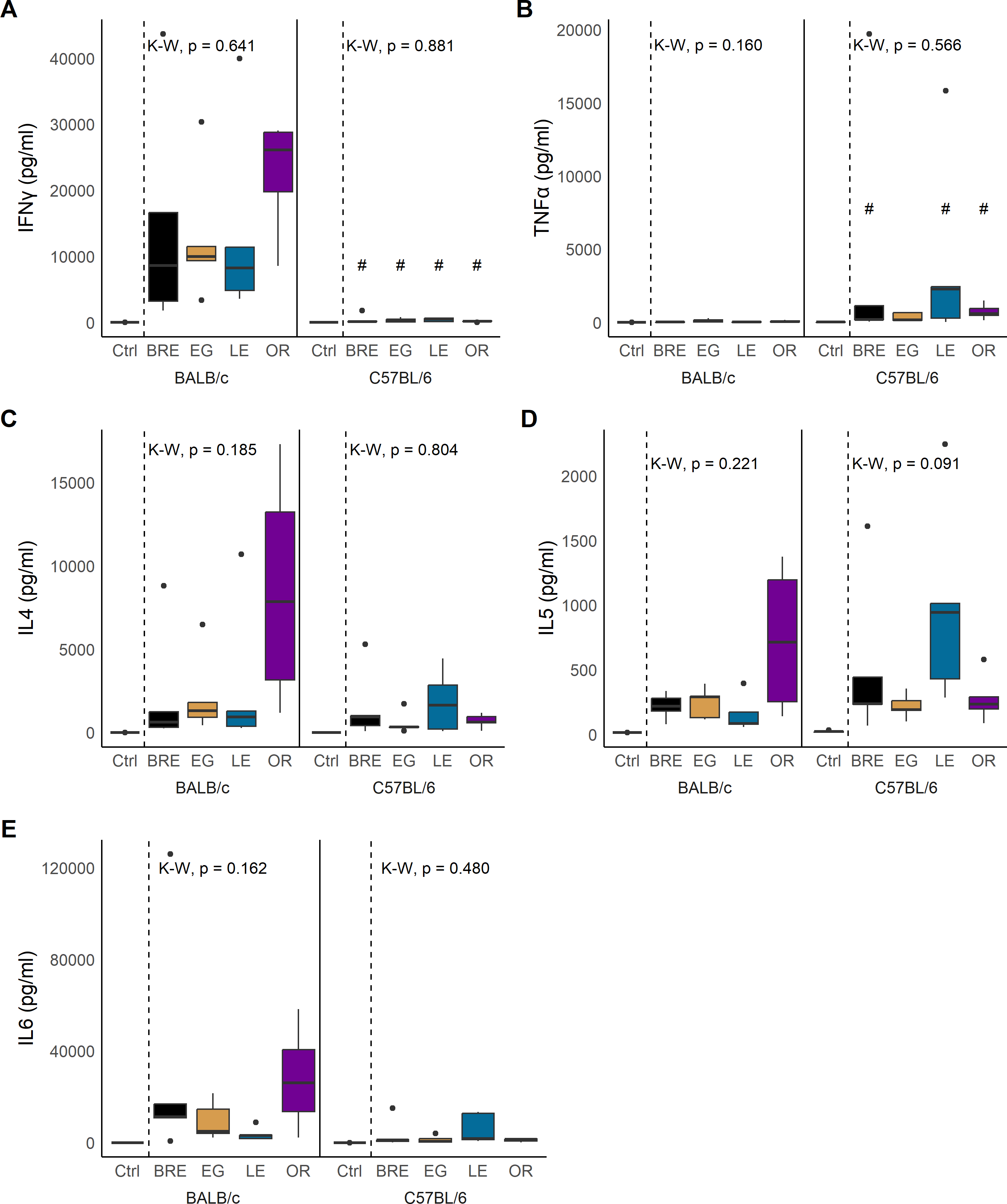
Cytokine production in response to schistosome infection. Box and whisker plots showing cytokine production of **(A)** IFN-γ, **(B)** TNF-α, **(C)** IL-4, **(D)** IL-5, and **(E)** IL-6 in BALB/c (left) and C57BL/6 mice (right) infected with the 4 parasite populations or uninfected controls. For all plots, statistical comparisons are between infected groups only using Kruskal-Wallis (K-W) followed by Dunn’s post hoc test. # *P* < 0.05, ## *P* < 0.01, ### *P* < 0.001: values are significantly different between host strains as calculated by Wilcoxon rank-sum test.

Host background did not significantly affect the production of cytokines IL-2, IL-4, IL-5, IL-6, IL-10, and IL-13 (Fig. 5C-E, Additional file 4: Fig. S4A-C), but IL-4, IL-5, and IL-6 exhibited substantial variation among all groups, with outcomes differing based on host-parasite interactions. For instance, SmOR infection led to an increase of IL-4 in BALB/c but not in C57BL/6 mice (Fig 5C; Wilcoxon; W = 13, *P* = 0.111). Conversely, SmLE infection resulted in almost significantly elevated levels of IL-5 in C57BL/6 but not in BALB/c mice (Fig. 5D; Wilcoxon; 5D: W = 1, *P* = 0.0636). Finally, IL-6 was higher in BALB/c mice infected with SmEG or SmOR compared to C57BL/6 mice (Fig. 5E; Wilcoxon; EG: W = 23, *P* = 0.0634; OR: W = 20, *P* = 0.0634) although this did not reach significance.

GLM analysis supported a significant impact of host strain on IFN-γ, TNF-α and IL-6 secretion (Table 4). We also revealed that IL-5 production was significantly impacted by host-parasite interactions. However, we observed no direct impact of parasite population on cytokine levels, which was consistent with the trait-by-trait analysis.

**Table 4.** Generalized linear model output for cytokine data.

| Characteristic | IFN-γ<sup>2</sup> | | | TNF-α<sup>2</sup> | | | IL2<sup>2</sup> | | | IL4<sup>2</sup> | | | IL5<sup>2</sup> | | |
| --- | --- | --- | --- | --- | --- | --- | --- | --- | --- | --- | --- | --- | --- | --- | --- |
|  | Beta | 95% CI<sup>1</sup> | p-value | Beta | 95% CI<sup>1</sup> | p-value | Beta | 95% CI<sup>1</sup> | p-value | Beta | 95% CI<sup>1</sup> | p-value | Beta | 95% CI<sup>1</sup> | p-value |
| Parasite |  |  | 0.6 |  |  | 0.11 |  |  | 0.085 |  |  | 0.2 |  |  | 0.2 |
| *BRE* | — | — |  | — | — |  | — | — |  | — | — |  | — | — |  |
| *EG* | 0.64 | -0.84, 2.1 |  | 2 | 0.14, 3.9 |  | 1 | -0.18, 2.3 |  | 1.2 | -0.57, 3.0 |  | 0.25 | -0.84, 1.3 |  |
| *LE* | 0.69 | -0.90, 2.3 |  | 0.35 | -1.7, 2.4 |  | -0.3 | -1.6, 1.1 |  | 0.8 | -1.1, 2.7 |  | -0.4 | -1.5, 0.82 |  |
| *OR* | 0.9 | -0.49, 2.3 |  | 1.3 | -0.44, 3.1 |  | 0.9 | -0.25, 2.1 |  | 1.8 | 0.15, 3.5 |  | 0.94 | -0.09, 2.0 |  |
| Host |  |  | <0.001* |  |  | 0.004* |  |  | 0.9 |  |  | 0.4 |  |  | >0.9 |
| *BALB/c* | — | — |  | — | — |  | — | — |  | — | — |  | — | — |  |
| *C57BL/6* | -4.7 | -6.0, -3.3 |  | 2.6 | 0.83, 4.4 |  | 0.1 | -1.1, 1.3 |  | -0.7 | -2.4, 0.96 |  | 0.03 | -1.0, 1.1 |  |
| Total eggs | 0 | 0.00, 0.00 | 0.11 | 0 | 0.00, 0.00 | 0.6 | 0 | 0.00, 0.00 | 0.7 | 0 | 0.00, 0.00 | 0.2 | 0 | 0.00, 0.00 | >0.9 |
| Parasite x Host |  |  | 0.3 |  |  | 0.2 |  |  | 0.2 |  |  | 0.4 |  |  | 0.005* |
| *EG x C57BL/6* | 1.1 | -0.90, 3.2 |  | -1.4 | -4.0, 1.2 |  | -0.1 | -1.7, 1.6 |  | -0.5 | -3.0, 2.0 |  | -0.2 | -1.7, 1.3 |  |
| *LE x C57BL/6* | 1.4 | -0.59, 3.4 |  | 1.6 | -0.93, 4.2 |  | 1.5 | -0.20, 3.1 |  | 0.85 | -1.6, 3.3 |  | 1.9 | 0.36, 3.3 |  |
| *OR x C57BL/6* | -0.3 | -2.4, 1.8 |  | -0.2 | -2.8, 2.4 |  | -0.2 | -1.9, 1.5 |  | -1.4 | -3.9, 1.1 |  | -0.9 | -2.4, 0.68 |  |

\* Statistically significant, <sup>1</sup> CI = Confidence Interval, <sup>2</sup> log-transformed dependent variable

| Characteristic | IL6<sup>2</sup> |  |  | IL10<sup>2</sup> |  |  | IL13<sup>2</sup> |  |  |
| --- | --- | --- | --- | --- | --- | --- | --- | --- | --- |
|  | Beta | 95% CI<sup>1</sup> | p-value | Beta | 95% CI<sup>1</sup> | p-value | Beta | 95% CI<sup>1</sup> | p-value |
| Parasite |  |  | 0.7 |  |  | 0.2 |  |  | 0.11 |
| *BRE* | — | — |  | — | — |  | — | — |  |
| *EG* | -0.1 | -2.0, 1.9 |  | 1.3 | -0.16, 2.7 |  | 1.6 | 0.27, 3.0 |  |
| *LE* | -0.9 | -2.9, 1.2 |  | -0.1 | -1.6, 1.4 |  | 1.4 | -0.12, 2.8 |  |
| *OR* | 0.38 | -1.4, 2.2 |  | 0.64 | -0.70, 2.0 |  | 0.89 | -0.40, 2.2 |  |
| Host |  |  | <0.001<sup>*</sup> |  |  | >0.9 |  |  | 0.2 |
| *BALB/c* | — | — |  | — | — |  | — | — |  |
| *C57BL/6* | -3.1 | -4.9, -1.3 |  | 0.05 | -1.3, 1.4 |  | 0.87 | -0.42, 2.2 |  |
| Total eggs | 0 | 0.00, 0.00 | 0.6 | 0 | 0.00, 0.00 | 0.8 | 0 | 0.00, 0.00 | 0.2 |
| Parasite x Host |  |  | 0.066 |  |  | 0.4 |  |  | 0.3 |
| *EG* x *C57BL/6* | 0.94 | -1.7, 3.5 |  | -1.4 | -3.3, 0.54 |  | -1.5 | -3.4, 0.34 |  |
| *LE* x *C57BL/6* | 3.2 | 0.61, 5.8 |  | 0.03 | -1.9, 2.0 |  | -1.2 | -3.0, 0.72 |  |
| *OR* x *C57BL/6* | 0.15 | -2.5, 2.8 |  | -1 | -3.0, 0.97 |  | -0.3 | -2.2, 1.7 |  |

<sup>\*</sup> Statistically significant, <sup>1</sup> CI = Confidence Interval, <sup>2</sup> log-transformed dependent variable

### Contribution of parasite population and host strain to infection phenotypes

Our trait-by-trait and GLM analyses suggest that parasitological traits tend to be determined by parasite genetics, while immunological phenotypes (CBC and cytokines) tend to be more affected by host genotype. We used effect size measures to directly partition the impact of parasite population, host genotype and host-parasite interactions on infection phenotypes. We categorized all phenotypes into parasitological, immunopathological, or immunological traits. We found that parasite population had a large and significant effect on three out of four parasitological traits, whereas host strain had a strong effect on seven of thirteen immunological traits (Fig. 6). Immunopathological traits were impacted by both host (five of six traits) and parasite (three of six traits). This analysis also identified significant host-parasite interaction effects for IL-5 secretion, consistent with the GLM analysis.

**Figure 6:**
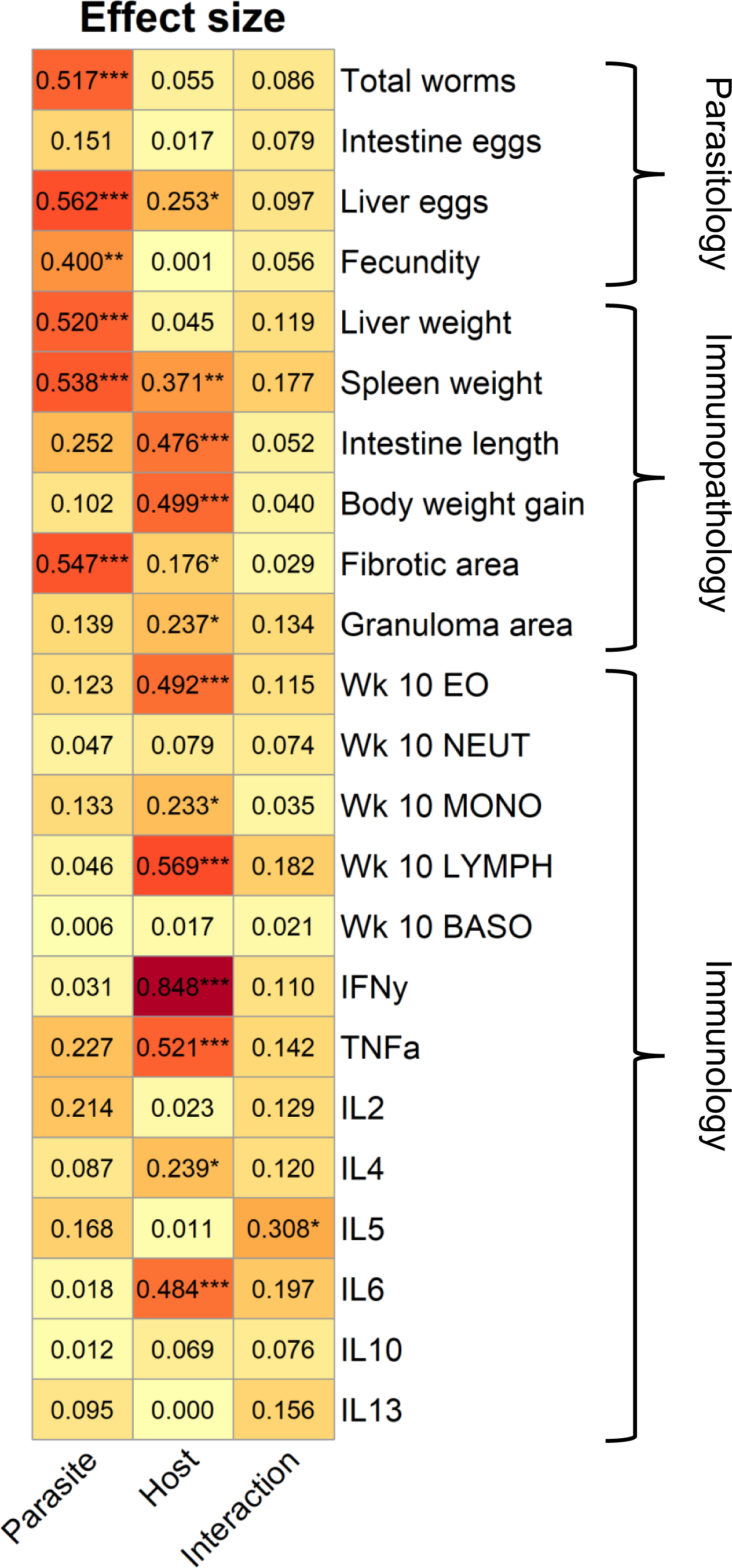
Effect Size by Parasite and Host Genotype on Disease Parameters. Heatmap showing effect size measures (Eta squared) as calculated by ANOVA. Parasite includes the 4 parasite populations, host includes both mouse strains, and interactions applies to the interaction between the two. Traits are organized in rows and assigned to the parasitology, immunopathology, or immunology category. * *P* < 0.05, ** *P* < 0.01, *** *P* < 0.001: values are significant per ANOVA result.

## Discussion

### Parasite and host genotype impact different disease parameters

We found that parasite and host genotype impact different categories of disease parameters during infection and post sacrifice. Out of four parasitological traits, parasite population strongly influenced the number of worms, liver egg burden and fecundity, whereas host influenced liver egg burden only. In contrast, host genotype impacted seven of 13 traits related to the immunological profile, with no discernible impact from the parasite population. In the immunopathology category (six traits), parasite genotype influenced three traits (liver weight, spleen weight, and fibrosis), while host genotype influenced five traits (spleen weight, intestine length, weight gain, fibrotic and granuloma area). Hence, the severity of immunopathology observed is dependent on both parasite and host.

### Parasite population effects on immunopathology

Egg burden is measured in epidemiological surveys because it is an important predictor of immunopathology in schistosomiasis [44–46]. Parasite genotype contributes to immunopathology in two ways in our experiments. First, we see significant differences between parasite populations in worm burden and fecundity. These jointly contribute to large differences in egg burden between parasite populations and impact liver and spleen weight and fibrosis. However, we also observe intrinsic differences between parasite lines, that impact immunopathology independent of egg burden. Our GLM analysis (Table 2) demonstrated that while egg burden contributed to variation in liver and spleen weight, parasite genotype also significantly impacted these traits. This is clearly shown by examining correlations between egg burden and organ weight (Additional file 5: Fig. S5).

This study confirms and extends our previous findings from studies of two Brazilian populations, SmBRE and SmLE, that show striking differences in cercarial shedding and sporocyst growth in the intermediate snail host [16,17,47]. These two parasite populations also showed the most extreme phenotype differences during infection in the vertebrate host. Despite comparable penetration success, we counted fewer worms and eggs and calculated lower fecundity in mice infected with SmBRE. Additionally, parameters such as liver weight, spleen weight, and fibrotic area were also lower in SmBRE-infected mice compared to the other schistosome populations characterized in the present study. We have previously demonstrated that fibrotic area and granuloma size were reduced in mice infected with parasite progeny from SmBRE and SmLE genetic crosses selected for low or high cercarial shedding from the aquatic snail [17]. While the experimental design in this study was rather different, we saw comparable differences in fibrotic area (lower in SmBRE than in SmLE), although granuloma size was unaffected. These observations underscore the profound impact of parasite factors on multiple phenotypes including immunopathology.

### Host genetic influences on susceptibility and immune profiles during infection

We did not detect differences in susceptibility and parasite fecundity between the mouse host strains. This supports findings from Alves et al. [41] who used SmLE and found no effect of mouse genetic background on worm and egg burden (Table 1). However, this is inconsistent with reports by Incani et al. [21], who documented higher worm but lower egg burden in BALB/c compared to C57BL/6 mice using Venezuelan and Brazilian *S. mansoni* populations, and Bin Dajem et al. [48] who found increased worm burden in C57BL/6 hosts with an Egyptian parasite. These differences could be explained by (i) the number of cercariae used for the infection (Alves: 30, Incani: 60, Bin Dajem: 100), (ii) the method of infection (tail immersion vs abdominal skin), (iii) statistical methods (log transformation vs non-parametric tests), (iv) the duration of the study, or (v) even genetic variation in the mouse colonies maintained at the different institutions [49].

IFN-γ, TNF-α, IL-6, monocyte and eosinophil levels were strongly influenced by host genotype as indicated by GLM and ANOVA analyses. This aligned with our expectations since C57BL/6 and BALB/c mice are known to induce different immune responses to many pathogens and are typically described as IFN-γ high/IL-4 low and IFN-γ low/IL-4 high responders respectively [22]. We observed comparable levels of IL-4 in BALB/c and C57BL/6 mice, but significantly increased IFN-γ (a known antifibrogenic agent) levels in infected BALB/c mice [50]. We hypothesize that this outcome represents hyperproduction in BALB/c mice, inducing protective immunity in response to schistosome infection [51,52]. This could prevent BALB/c mice from mounting a potent type 2 immune response, impact granuloma formation, and explain the mortality we observed in BALB/c and not in C57BL/6 mice [53–55].

We observed larger granulomas, which are predominantly driven by a type 2 immune response, in C57BL/6 mice [40,43,56]. In contrast, Alves et al. [41] measured larger granulomas in BALB/c mice. In addition to high IFN-γ levels, we also saw significantly more eosinophils at week 10 post infection in the blood of BALB/c mice compared to C57BL/6 mice. This could imply that eosinophils in C57BL/6 mice were recruited to the liver tissue, because granulomas surrounding *S. mansoni* eggs are mainly composed of eosinophils [56,57]. However, that eosinophil depletion did not change granuloma size in schistosome infected mice contradicts this idea [58]. Future studies could follow up on this observation using blood and whole spleen flow cytometry to distinguish the different immune cell populations in both *S. mansoni* infected mouse hosts.

Both GLM and ANOVA revealed a significant impact of host-parasite interactions on IL-5 secretion. IL-5 influences various cell types including B cells and eosinophils in a pleiotropic manner [59,60]. This suggests that the four schistosome populations may differ in antigenicity, resulting in varying levels of induced inflammation affected by mouse host genetics. The fact that IL-5 is involved in fibrosis regulation, coupled with our analyses revealing a nearly significant impact of parasite population on fibrosis, supports this notion [61].

### Limitations of this study

The influence of parasite genetics on immunopathological parameters measured in this study is likely to be conservative. Laboratory schistosome populations maintain surprisingly high levels of genetic and phenotypic variation [62, Jutzeler et al., in prep]. By infecting mice with genetically diverse schistosome larvae from four parasite populations, we captured an average phenotype for each population, so extreme phenotype traits will be masked. We expect that use of genetically homogeneous parasite populations would result in much larger effects of parasite genotype. Future studies could focus on phenotype measures in mice infected with inbred schistosome lines generated by serial inbreeding over several generations. Such inbred schistosome lines are not currently available, but could provide a valuable resource for investigating the role of parasite genetics in determining outcome of infection.

Similarly, we probably overestimated the role of host genetics by conducting these experiments using two inbred mouse strains known to show divergent immune responses to infection. Future work on host influences on schistosome immunopathology could use inbred mouse lines from the collaborative cross to determine the host genes involved [63].

## Conclusions

This study highlights the significant influence of both schistosome parasite and mouse host genetics on immunopathological parameters. We found that (i) parasite population influenced liver and spleen weight and fibrotic area, (ii) both differences in total egg burden between parasite populations and intrinsic parasite factors unrelated to egg burden contribute to immunopathology, and (ii) there were significant host-parasite interactions on IL-5 production. We anticipate that the impact of parasite genotype on immunopathology will be more pronounced when using inbred parasite lines; such inbred lines could also simplify genetic analysis of immunopathology traits.

## Supplementary information

**Additional file 1: Figure S1.** Box plot showing the ratio of liver eggs to intestine eggs in schistosome infected BALB/c (left) and C57BL/6 mice (right). Statistical comparison between parasite populations done for each host with Kruskal-Wallis (K-W) or ANOVA (BALB/c: Kruskal-Wallis; H = 4.26, df = 3, *P* = 0.235; C57BL/6: ANOVA; F_(3, 12)_ = 1.34, *P* = 0.306).

**Additional file 2: Figure S2.** Intestine length of schistosome infected BALB/c and C57BL/6 mice. Box plot showing intestine length in BALB/c (left) and C57BL/6 mice (right) infected with the 4 parasite populations or uninfected controls. ANOVA to identify comparisons between mice (BALB/c: F_(3, 16)_ = 1.28, *P* = 0.323; C57BL/6: F_(3, 16)_ = 3.19, *P* = 0.052) and T-test to compare hosts (*t*_(5.89)_ = 6.10, *P* = 0.00378). # *P* < 0.05, ## *P* < 0.01, ### *P* < 0.001: values are significantly different between host strains.

**Additional file 3: Figure S3.** Neutrophil and reticulocyte levels and mean hematocrit (HCT) output. Longitudinal plots showing select CBC levels in BALB/c (top) and C57BL/6 mice (bottom) infected with the 4 parasite populations (solid lines) or uninfected control (dashed lines). **(A)** Neutrophil levels (Wilcoxon test to compare hosts: Baseline: W = 264, *P* = 0.0423; Week 4: W = 275, *P* = 0.0344). **(B)** Hematocrit (HCT) (Friedman; BALB/c: χ2 = 1.68, df = 3, *P =* 0.642; C57BL/6: χ2 = 3.8, df = 3, *P =* 0.284). Plot shows significant differences between hosts at baseline, and weeks 4, 6, and 8 (Wilcoxon test; Baseline: W = 101, *P* = 0.0324; Week 4: W = 84, *P* = 0.0156; Week 6: W = 77, *P* = 0.0156; Week 8: W = 94.5, *P* = 0.0258). **(C)** Reticulocyte production (Friedman; BALB/c: χ2 = 5.8, df = 3, *P =* 0.122; C57BL/6: χ2 = 6.6, df = 3, *P =* 0.086). # *P* < 0.05, ## *P* < 0.01, ### *P* < 0.001: values are significantly different between host strains.

**Additional file 4: Figure S4:** No differences in IL-2, IL-10, and IL-13 levels. **(A)** IL-2 (Kruskal-Wallis; BALB/c: H = 7.34, df = 3, *P* = 0.0618; C57BL/6: H = 0.509, df = 3, *P* = 0.917), **(B)** IL-10 (Kruskal-Wallis; BALB/c: H = 2.87, df = 3, *P* = 0.411; C57BL/6: H = 2.64, df = 3, *P* = 0.451), and **(C)** IL-13 (Kruskal-Wallis; BALB/c: H = 1.13, df = 3, *P* = 0.771; C57BL/6: H = 1.94, df = 3, *P* = 0.584) levels were not significantly different between parasite populations or mouse lines. K-W = Kruskal-Wallis.

**Additional file 5: Figure S5:** Correlation plots between total egg counts and liver/spleen weight. **(A)** Normalized liver weight is significantly correlated in two out of four parasite populations (Spearman’s correlation coefficient; BRE: *rs* = 0.85, *P* = 0.006, EG: *rs* = 0.24, *P* = 0.582, LE: *rs* = 0.71, *P* = 0.058, OR: *rs* = 0.90, *P* = 0.002). **(B)** Only SmBRE egg counts are significantly correlated with total egg burden (Spearman’s correlation coefficient; BRE: *rs* = 0.75, *P* = 0.025, EG: *rs* = 0.17, *P* = 0.703, LE: *rs* = 0.48, *P* = 0.243, OR: *rs* = 0.53, *P* = 0.148).

**Additional file 6: Table S1:** Zenodo repository information for each histology image data set.

## Declarations

### Acknowledgements

We thank the animal care staff of the Southwest National Primate Research Center (SNPRC) for providing rodent care, training and equipment to perform this study; Renee Escalona, Colin Chuba and Jesse Martinez from the Hixon Pathology core lab (Texas Biomedical Research Institute) for performing all the histopathology preparations and staining of mouse livers and Dr. Vinay Shivanna for providing access to the HALO software for analyzing the histopathology slides; and Dr. Vida Hodara from the Biology core lab for processing the cytokine samples. Finally, we thank Dr. P’ng Loke, Chief of the Type 2 Immunity Section at the NIAID and Dr. Reagan Meredith (Texas Biomedical Research Institute) for their input and recommendations regarding study design and statistical analysis respectively.

### Funding

This research was supported by a Graduate Research in Immunology Program training grant NIH T32 AI138944 (KSJ), and NIH R21 AI171601-02 (FDC, WL), R01 AI133749, R01 AI166049 (TJCA), and was conducted in facilities constructed with support from Research Facilities Improvement Program grant C06 RR013556 from the National Center for Research Resources. SNPRC research at Texas Biomedical Research Institute is supported by grant P51 OD011133 from the Office of Research Infrastructure Programs, NIH.

### Availability of data and materials

The datasets supporting the conclusions of this article and all codes used for data analysis and generation of figures (2-6, S1-S5) are available at https://github.com/kathrinsjutzeler/sm_immunopathology/ and https://doi.org/10.5281/zenodo.10465595. All histology images used in this manuscript were uploaded to Zenodo. Additional file 6: Table S1 contains the description and repository information for each data set.

### Author contributions

KSJ and TJCA designed and planned the experiments. WL and FDC provided training and assisted with methodology. KSJ performed all the experiments (mouse infections, measurement of all the phenotypes in hosts, and cytokine assay), and all the data analyses including liver histopathology. KSJ and TJCA drafted the manuscript. All authors read and approved the final manuscript.

### Competing interests

The authors declare that they have no competing interests.

## Notes

### Competing Interest Statement

The authors have declared no competing interest.

