## Supplementary figures and images for "Contribution of parasite and host genotype to immunopathology of schistosome infections"

### Supplemental Figure 1

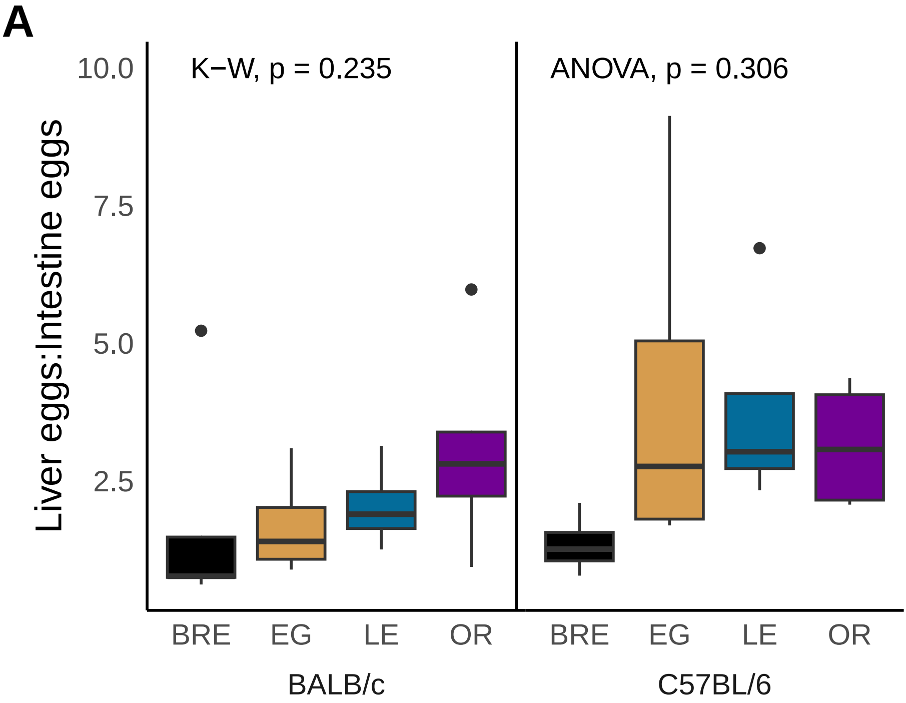

### Supplemental Figure 2

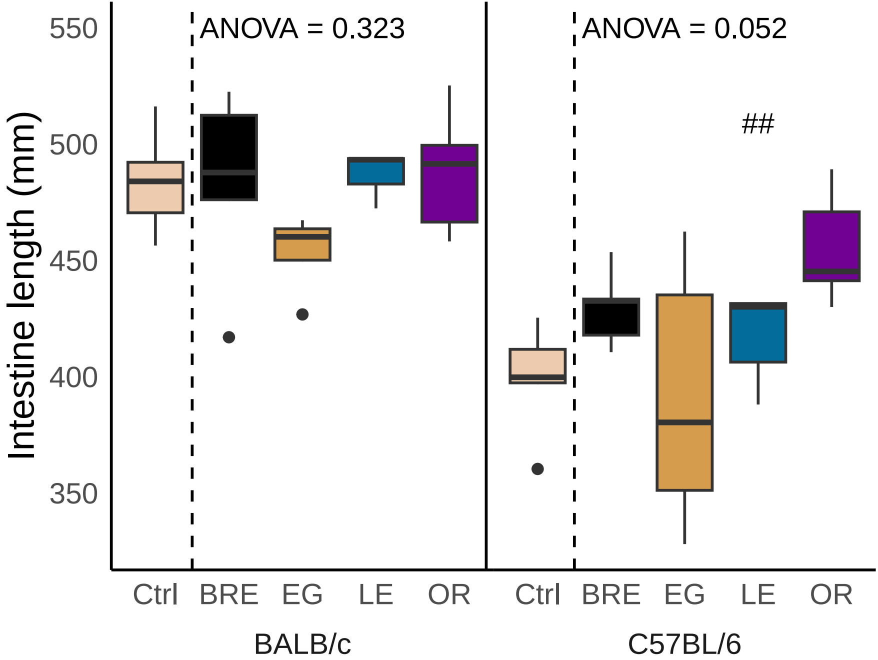

### Supplemental Figure 3

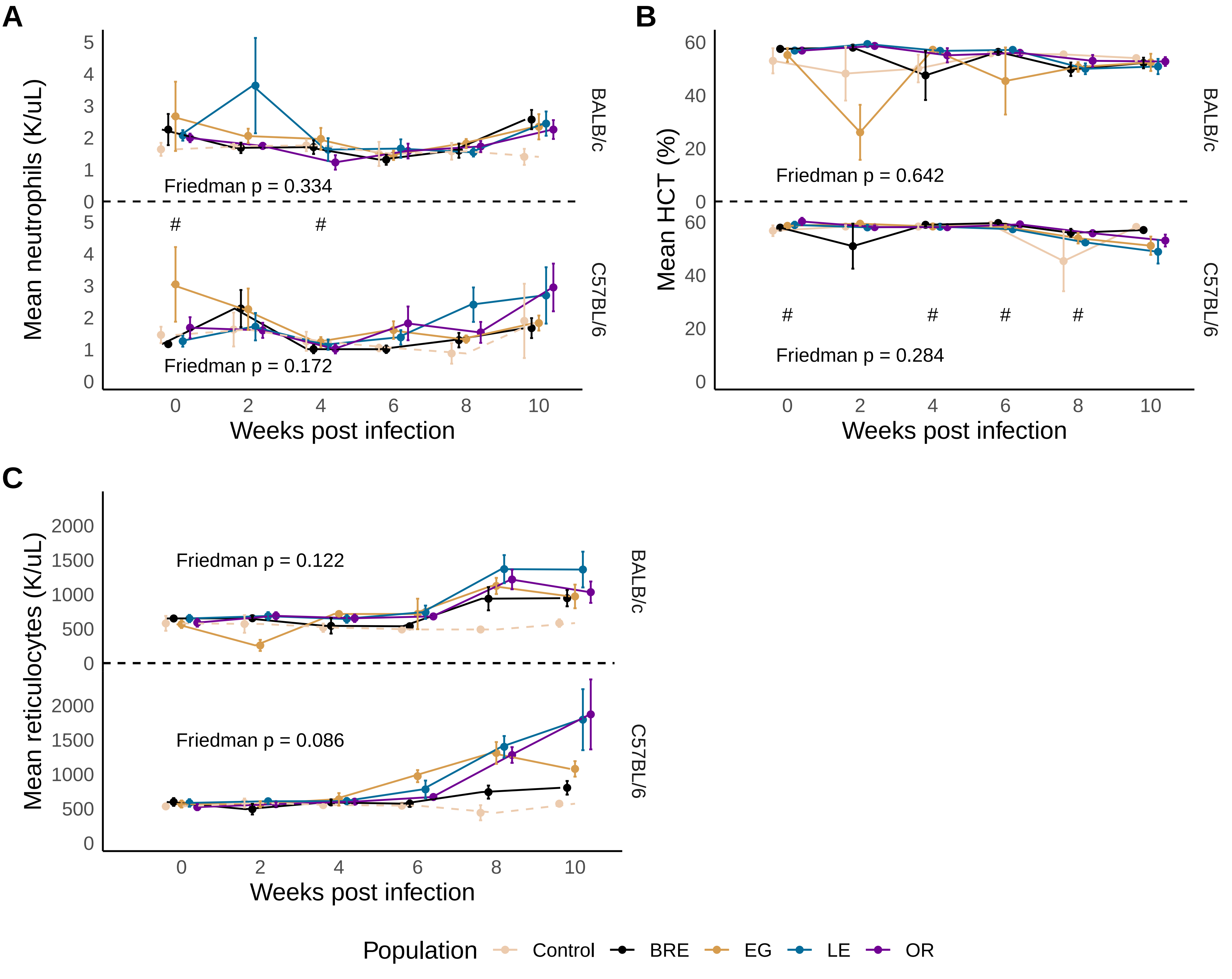

### Supplemental Figure 4

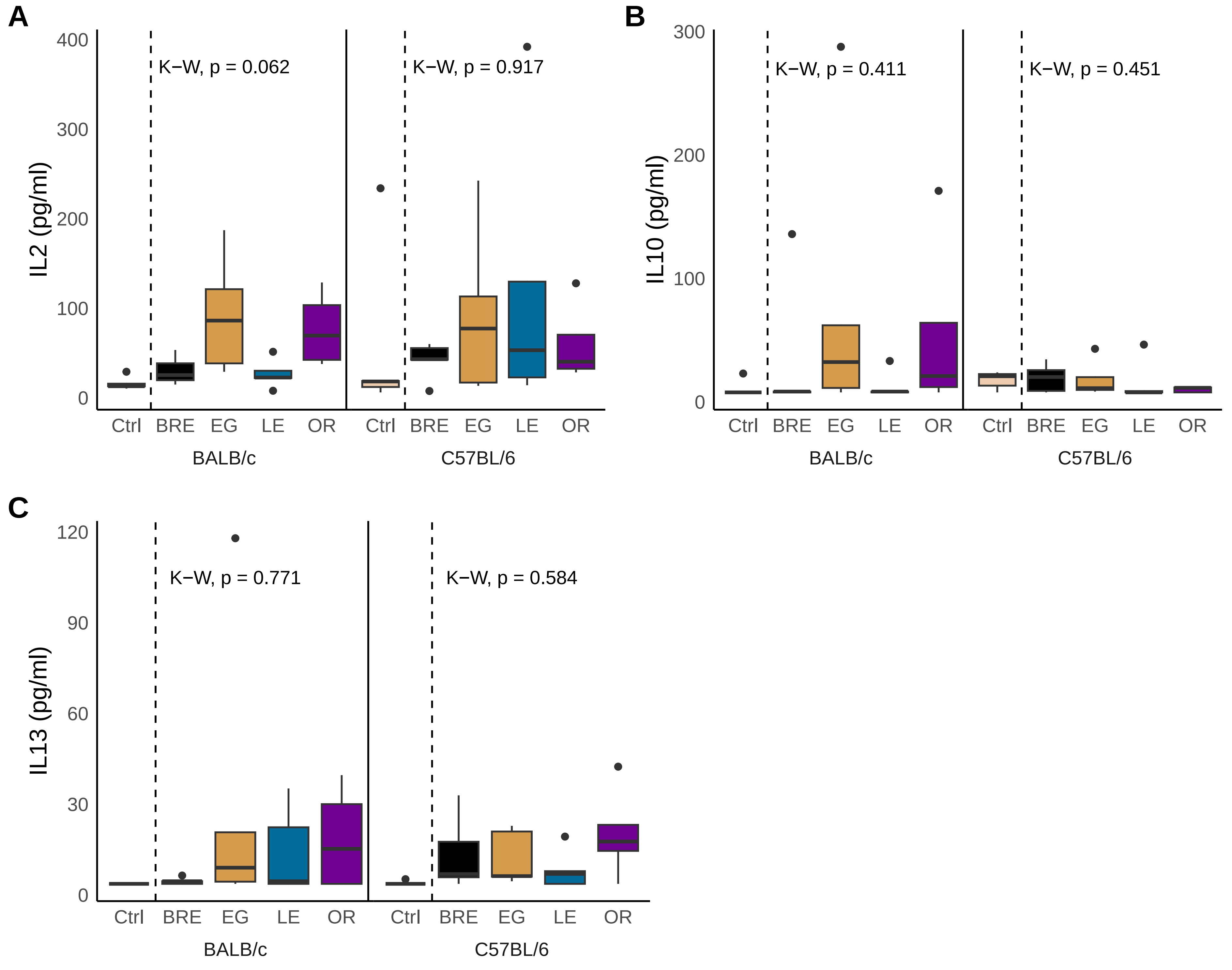

### Supplemental Figure 5

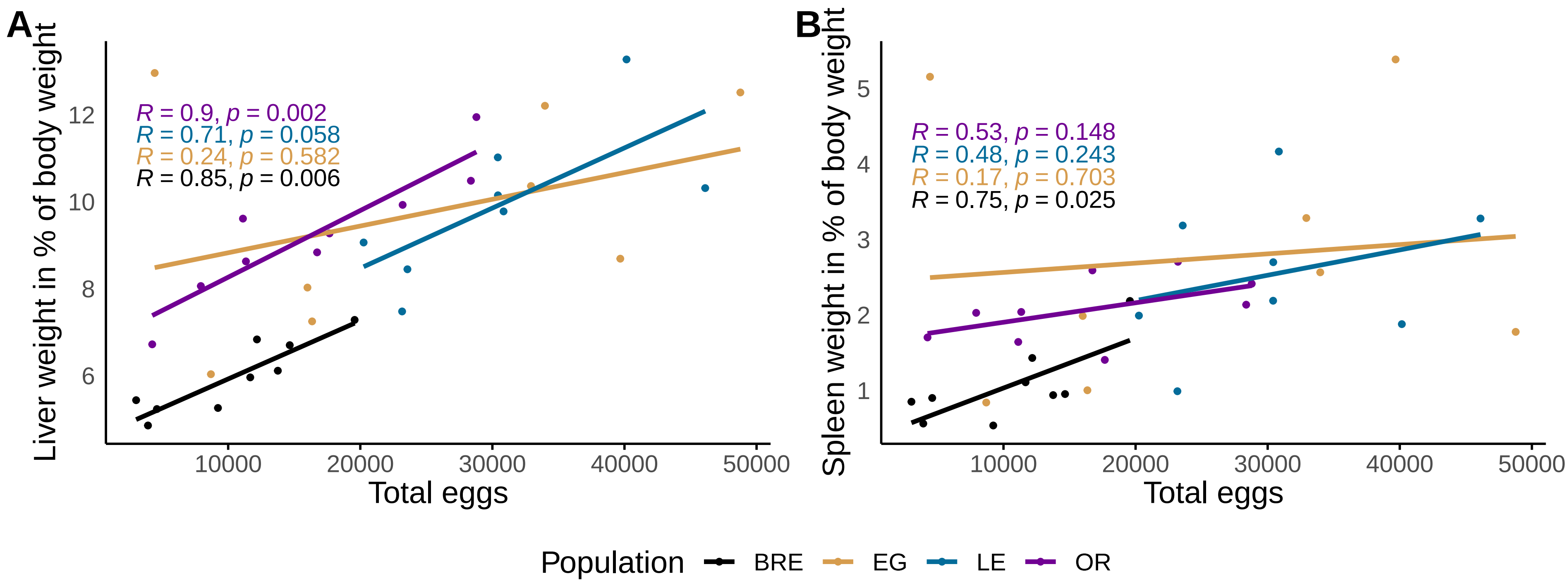
