## Supplemental Table 1 for "Contribution of parasite and host genotype to immunopathology of schistosome infections"

| Additional file 5: Table S1. Zenodo repository information for histology images | | |
| --- | --- | --- |
| Name | Identifier | DOI |
| BALB/c_BRE_H&E_images | 10162056 | 10.5281/zenodo.10162056 |
| BALB/c_BRE_Trichrome_images | 10162078 | 10.5281/zenodo.10162078 |
| BALB/c_control_H&E_images | 10161988 | 10.5281/zenodo.10161988 |
| BALB/c_control_Trichrome_images | 10161996 | 10.5281/zenodo.10161996 |
| BALB/c_EG_H&E_images | 10162062 | 10.5281/zenodo.10162062 |
| BALB/c_EG_Trichrome_images | 10162082 | 10.5281/zenodo.10162082 |
| BALB/c_LE_H&E_images | 10162066 | 10.5281/zenodo.10162066 |
| BALB/c_LE_Trichrome_images | 10162088 | 10.5281/zenodo.10162088 |
| BALB/c_OR_H&E_images | 10162074 | 10.5281/zenodo.10162074 |
| BALB/c_OR_Trichrome_images 1/2 | 10162092 | 10.5281/zenodo.10162092 |
| BALB/c_OR_Trichrome_images 2/2 | 10235215 | 10.5281/zenodo.10235215 |
| C57BL/6_BRE_H&E_images | 10162012 | 10.5281/zenodo.10162012 |
| C57BL/6_BRE_Trichrome_images | 10162034 | 10.5281/zenodo.10162034 |
| C57BL/6_control_H&E_images | 10044960 | 10.5281/zenodo.10044960 |
| C57BL/6_control_Trichrome_images | 10161827 | 10.5281/zenodo.10161827 |
| C57BL/6_EG_H&E_images 1/2 | 10162018 | 10.5281/zenodo.10162018 |
| C57BL/6_EG_H&E_images 2/2 | 10234748 | 10.5281/zenodo.10234748 |
| C57BL/6_EG_Trichrome_images 1/2 | 10162038 | 10.5281/zenodo.10162038 |
| C57BL/6_EG_Trichrome_images 2/2 | 10234782 | 10.5281/zenodo.10234782 |
| C57BL/6_LE_H&E_images 1/2 | 10162028 | 10.5281/zenodo.10162028 |
| C57BL/6_LE_H&E_images 2/2 | 10234790 | 10.5281/zenodo.10234790 |
| C57BL/6_LE_Trichrome_images 1/2 | 10162042 | 10.5281/zenodo.10162042 |
| C57BL/6_LE_Trichrome_images 2/2 | 10234800 | 10.5281/zenodo.10234800 |
| C57BL/6_OR_H&E_images 1/2 | 10162030 | 10.5281/zenodo.10162030 |
| C57BL/6_OR_H&E_images 2/2 | 10234716 | 10.5281/zenodo.10234716 |
| C57BL/6_OR_Trichrome_images 1/2 | 10162044 | 10.5281/zenodo.10162044 |
| C57BL/6_OR_Trichrome_images 2/2 | 10234814 | 10.5281/zenodo.10234814 |
